# Distinct Virome and Bacteriome Profiles of Term and Preterm Placentas from African, Asian and European Women

**DOI:** 10.64898/2025.12.11.693726

**Authors:** Khondoker M Akram, Mustafizur Rahman, Tahsin Khan, Emmanuel Amabebe, Nadia Ikumi, Muntasir Alam, Shafina Jahan, Ally Oosthuizen, Cara Mahon, Skye Barker, Helen Hipperson, Marta Cohen, Mushi Matjila, Dilly OC Anumba

**Affiliations:** Division of Clinical Medicine, University of Sheffield, Sheffield, United Kingdom; Infectious Diseases Division, icddr,b, Dhaka, Bangladesh; Department of Obstetrics and Gynaecology, University of Texas Medical Branch at Galveston, Galveston, Texas, USA; Department of Obstetrics and Gynaecology, University of Cape Town, Cape Town, South Africa; School of Biosciences, University of Sheffield, Sheffield, United Kingdom; Sheffield Children’s NHS Foundation Trust, Sheffield, United Kingdom

**Keywords:** Placental virome and bacteriome, preterm birth, shotgun metagenomic study

## Abstract

While the human placental microbiome has been extensively studied, information regarding the virome remains limited. The association between subclinical placental viral infections and spontaneous preterm birth (PTB) is poorly understood. Hence, we examined fresh placenta samples from women in the UK, Bangladesh, and South Africa to elucidate the virome profiles and association with spontaneous PTB. We employed shotgun metagenomics, real-time PCR, and Gram staining, complemented by bioinformatics, to detect both viruses and bacteria in the decidua and chorionic villous tissues. We did not identify any known pathogenic viruses in 236 placental samples, except for one preterm placenta from the Bangladeshi cohort, where human herpesvirus 6 was detected. The majority of the viruses were bacteriophages present at very low abundance and low frequencies across the three cohorts. *Lambdavirus lambda* was detected in 14% of UK and 11% of Bangladeshi samples but was absent in South African samples. *Papiine betaherpesvirus 3* was identified in 62.5% of South African placentas. *Punavirus* (29.1%) and Streptococcus phages (16.67%) were also prevalent in South African samples. A variety of bacterial species including *Moraxella osloensis*, *Delftia lacustris*, *Cutibacterium acnes, Escherichia coli* and *Staphylococcus epidermidis* were detected in samples across the three cohorts in variable frequencies but without any notable tissue immune responses. None of the viruses or bacteria exhibited associations with PTB, except *D. lacustris*, which was significantly more abundant in preterm compared to term placentas in the Bangladeshi cohort. We conclude that the human placenta harbours a virome predominated by bacteriophages and that subclinical viral infection in spontaneous PTB is rare.

## 1. MAIN

The existence of a distinct placental microbiome has been the subject of intense debate in recent years. The human placenta was traditionally believed to be sterile in healthy conditions^1^. This view was challenged by the detection of bacteria in the basal plates of term and preterm placentas in the absence of chorioamnionitis^2, 3^. Subsequently, a unique placental microbiome, composed of non-pathogenic commensal species—including *Escherichia coli*, *Bacteroides* spp., *Staphylococcus epidermidis*, and *Propionibacterium acnes*—was identified by 16S ribosomal DNA sequencing and whole-genome shotgun metagenomic methods^4^. Furthermore, a recent study demonstrated the presence of low-abundance, low-biomass microbial communities within the chorionic villi of term and preterm placentas in the absence of clinical or histologic chorioamnionitis^5^.

In contrast, another study utilising rigorous contamination controls on human placenta samples found no evidence of a microbiome in most placentas^6^. The researchers noted that nearly all microbial signals detected were attributable to contamination during labour and delivery or bacterial DNA contamination in reagents. However, the study identified *Streptococcus agalactiae* in 5% of the samples as a genuine infection. Subsequent studies also failed to confidently identify any discernible microbiome in the placentas^1, 7^.

The potential contamination of reagents in sequence-based metagenomic analyses presents an inherent technical challenge, particularly when analysing low-biomass samples such as the placenta^8, 9^. Nonetheless, the existence of a microbiota continuum along the female reproductive tract - comprising the cervicovaginal canal, uterus, and fallopian tubes - along with the hypervascularity of the villous placenta, indicates a potential for ascending or haematogenous spread of microorganisms to the placenta^10, 11^. While most studies have focused on detecting bacterial microbiomes in the placenta through DNA and 16S rRNA amplification techniques, a recent investigation employed mass spectrometry to identify viral proteins primarily associated with various bacteriophages in placentas^12^. This study concluded that changes in placental virome diversity might contribute to the pathogenesis of fetal growth restriction (FGR).

The placental virome remains understudied but consists of both endogenous retroviral elements integrated into the host genome and transient exogenous viral infections acquired during pregnancy. Exogenous viral infections such as cytomegalovirus (CMV), herpes simplex virus (HSV), human papillomavirus (HPV), human parvovirus B19, and SARS-CoV-2 are well-documented contributors to adverse pregnancy outcomes, including preterm birth (PTB), defined as delivery before 37 weeks of gestation^13, 14, 15, 16, 17, 18^. However, our understanding of the placental virome and its role in both normal and adverse pregnancy outcomes remains limited and contentious. Furthermore, the significance of subclinical viral infections of the placenta in spontaneous PTB is poorly understood. Studies have identified viral DNA and RNA in placental tissues linked to PTB, even in the absence of maternal symptoms^19, 20^. Research indicates that viral infections can impair trophoblast function and trigger a localised inflammatory cascade within the placenta, ultimately resulting in the premature activation of parturition pathways associated with PTB^21, 22, 23, 24^.

In this study, our primary aims were: (i) to identify virome profiles in placentas from diverse racial and geographical backgrounds, and (ii) to investigate whether subclinical viral infections in the placenta are associated with spontaneous PTB. To address the inherent challenges of reagent- and sequencing-related contamination, we collected fresh placentas from three geographical regions: the UK, Bangladesh, and South Africa, and extracted total RNA and DNA from tissue biopsies using the same reagents in a sterile laboratory environment in the UK. The extracted nucleic acids were then analysed via shotgun metagenomic sequencing using the same GridION sequencer. This standardised approach enabled us to discern reagent-associated contaminants in our metagenomic data. Additionally, DNA was extracted from cervicovaginal fluid (CVF) samples and placental tissues from the same women who delivered vaginally to differentiate placental virome/bacteriome signals from potential vaginal microbiota contamination. To address technique-specific sensitivity issues, we conducted a real-time PCR array on a subset of placental samples, targeting eight commonly occurring pathogens. Finally, Gram staining was performed on a subset of formalin-fixed paraffin-embedded (FFPE) placental samples to locate bacterial presence.

Our shotgun metagenomic sequencing and real-time PCR array did not identify any known pathogenic viruses in 236 placental samples, except for one preterm placenta from the Bangladeshi cohort, where human herpesvirus 6 (HHV-6) was detected using the PCR method. Among other viral species, the majority were bacteriophages, present at very low abundance and low frequencies in placental samples across the three countries. A variety of bacterial species were also detected in placental samples across the three countries.

## 2. RESULTS

### 2.1. Participant criteria and sampling

During the period of 2019 to 2021, a total of 340 fresh placentas were collected from pregnant women (age > 16 years) who delivered spontaneously at term (> 37 weeks of gestation) or preterm (< 37 weeks of gestation) in our PRIME preterm birth research consortium partner countries: the UK (n = 107), Bangladesh (n = 176), and South Africa (n = 57). The villous/decidua tissue samples from 236 placentas were utilised for shotgun metagenomic and real-time PCR array analyses. Villous/decidua (50:50) tissue samples from 96 placentas underwent shotgun metagenomic sequencing (UK: term = 18, preterm = 18; Bangladesh: term = 18, preterm = 18; South Africa: term = 14, preterm = 10). In addition, villous/decidua tissues from 175 Bangladeshi placentas (term = 119, preterm = 56) were used for a real-time PCR array analysis targeting eight commonly occurring pathogens associated with adverse pregnancy outcomes, including HSV-1, HSV-2, hCMV, HHV-6, HHV-7, B-19, *Enterovirus,* and *Treponema pallidum* (Figure 1, Table S1).

**Figure 1:**
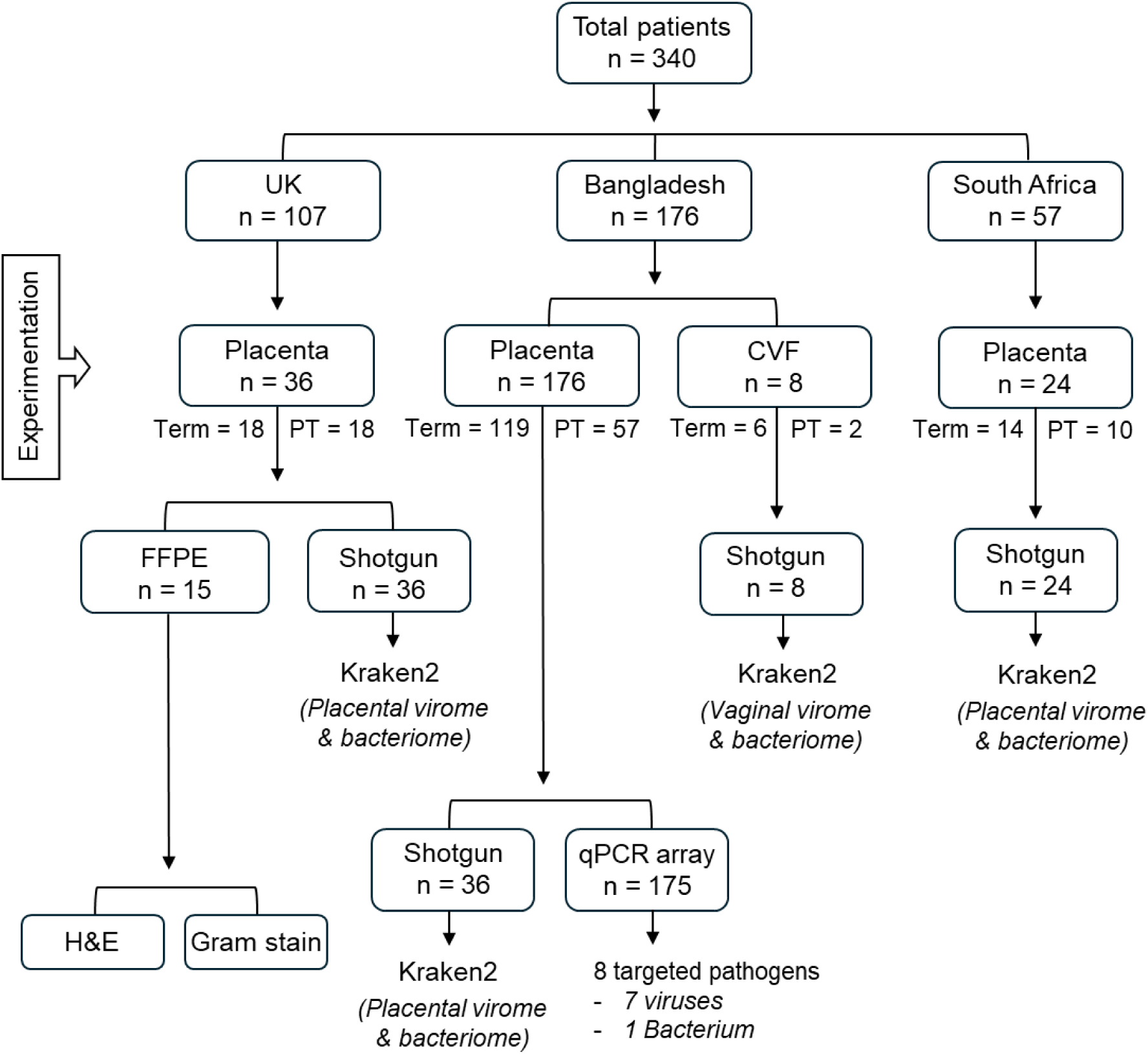
Flowchart illustrating the stratification of the patient’s samples and the methods of experimentation. CVF, cervicovaginal fluid; FFPE, formalin-fixed paraffin-embedded; H&E, haematoxylin and eosin staining; PT, preterm; qPCR, quantitative polymerase chain reaction.

Shotgun metagenomic sequencing was also conducted on cervicovaginal fluid (CVF) samples obtained by mid-vaginal swabs from the Bangladeshi pregnant women at 22-24 weeks of gestation who spontaneously delivered vaginally (n = 8) either at term (n = 6) or preterm (n = 2). These CVF samples were collected from the women whose placentas were included for metagenomic sequencing as well (hence called linked samples). Metagenomic data from these matched placenta/CVF samples were used to assess potential contamination of placenta by the vaginal microbial species.

Women with singleton pregnancies who delivered spontaneously, whether at term or preterm, vaginally or via caesarean section, were included in this study. Women exhibiting symptoms of genitourinary tract or systemic infections were excluded from the study across the participating country sites. The participants from the UK were predominantly White, while those in South Africa were primarily Black. Participants in Bangladesh were solely Asian Bangladeshi. Detailed recruitment criteria and demographic characteristics of participants are outlined in Table 1.

**Table 1:** Demographic characteristics of the participants. Values are presented as means with standard deviations (±SD), or ‘n’ numbers with percentages in the parentheses. p values were calculated by either Mann-Whitney U test or ^#^Fisher’s exact test between term and preterm groups. Maternal body mass indices (BMI) of the South African cohort were not recorded.

|  | UK<br>(n = 36) |  | P<br>values | Bangladesh<br>(n = 176) |  | P<br>values | South Africa<br>(n = 24) |  | P<br>values |
| --- | --- | --- | --- | --- | --- | --- | --- | --- | --- |
|  | Term<br>(n = 18) | Preterm<br>(n = 18) |  | Term<br>(n = 119) | Preterm<br>(n = 57) |  | Term<br>(n = 14) | Preterm<br>(n = 10) |  |
| Maternal age, years<br>(±SD) | 31.51<br>(5.27) | 27.58<br>(6.92) | 0.0954 | 24.94<br>(4.39) | 23.93<br>(5.8) | 0.0504 | 32.45<br>(5.27) | 31.55<br>(7.11) | 0.8523 |
| #Maternal race, n (%) |  |  |  |  |  |  |  |  |  |
| White | 16<br>(88.8) | 17<br>(94.4) | >0.9999 | 0 (0) | 0 (0) | >0.9999 | 0 (0) | 1 (10) | 0.4167 |
| Black | 1 (5.6) | 0 (0) |  | 0 (0) | 0 (0) |  | 14 (100) | 9 (90) |  |
| Asian and others | 1 (5.6) | 1 (5.6) |  | 119<br>(100) | 57 (100) |  | 0 (0) | 0 (0) |  |
| Gestational age, weeks<br>(±SD) | 39.25<br>(1.15) | 33.03<br>(3.8) | <0.0001 | 39.02<br>(1.66) | 34.63<br>(2.57) | <0.0001 | 38.64<br>(0.72) | 34.6<br>(1.85) | <0.0001 |
| #Mode of delivery, n (%) |  |  |  |  |  |  |  |  |  |
| Vaginal | 17<br>(94.4) | 15<br>(83.3) | 0.6026 | 81<br>(68.1) | 47<br>(82.5) | 0.0484 | 4 (28.6) | 8 (80) | 0.0361 |
| Caesarean | 1 (5.6) | 3 (16.7) |  | 38<br>(31.9) | 10<br>(17.5) |  | 10 (71.4) | 2 (20) |  |
| #Membrane rupture, n<br>(%) |  |  |  |  |  |  |  |  |  |
| SROM | 12<br>(66.7) | 1 (5.6) | <0.0001 | 52<br>(43.7) | 46<br>(80.7) | <0.0001 | 7 (50) | 8 (89) | 0.3875 |
| PPROM | 0 (0) | 8 (44.4) |  | 2 (1.7) | 0 (0) |  | 1 (7.1) | 0 (0) |  |
| AROM | 6 (33.3) | 9 (50) |  | 65<br>(54.6) | 11<br>(19.3) |  | 6 (42.9) | 2 (20) |  |
| BMI at delivery, kg/m2<br>(±SD) | 28.27<br>(6.64) | 27.27<br>(8.32) | 0.6591 | 27.59<br>(3.8) | 25.63<br>(4.11) | 0.0002 | NA | NA | NA |
| #Diabetes mellitus, n (%) |  |  |  |  |  |  |  |  |  |
| Yes | 0 (0) | 0 (0) | >0.9999 | 0 (0) | 1 (1.8) | 0.3239 | 0 | 1 (10) | 0.4167 |
| No | 18 (100) | 18 (100) |  | 119<br>(100) | 56<br>(98.2) |  | 14 (100) | 9 (90) |  |
| #Smoking, n (%) |  |  |  |  |  |  |  |  |  |
| Yes | 3 (16.7) | 4 (22.2) | >0.9999 | 0 (0) | 0 (0) | >0.9999 | 0 (0) | 1 (10) | 0.4167 |
| No | 15<br>(83.3) | 14<br>(77.8) |  | 119<br>(100) | 57 (100) |  | 14 (100) | 9 (90) |  |
| Newborn weight, kg<br>(±SD) | 3.4 (0.4) | 2 (0.7) | <0.0001 | 2.8<br>(0.41) | 2.27<br>(0.56) | <0.0001 | 3.3 (0.4) | 2.3 (0.8) | 0.0031 |
| Placenta weight, grams<br>(±SD) | 691.1<br>(121) | 446.5<br>(141.9) | <0.0001 | 478.5<br>(90) | 400.8<br>(140) | <0.0001 | 632.9<br>(114.5) | 507<br>(181.1) | 0.0537 |

**Note:** SROM: spontaneous rupture of membranes; PPRM: preterm premature rupture of the membranes; AROM: artificial rupture of membranes; NA: data not available.

### 2.2. Distinct virome profiles are detected in term and preterm placentas delivered by the UK, Bangladeshi and South African women

Our shotgun metagenomic sequencing of 96 chorionic villous/decidua basalis tissue samples (50:50) from fresh placentas delivered in the UK, Bangladesh, and South Africa did not detect any known pathogenic DNA or RNA viruses. However, we identified the *BeAn 58058 virus* (BAV), a variant of the Vaccinia virus, in the majority of placenta samples across the countries, with prevalences of 52.2%, 88.89%, and 100% in the samples from the UK, Bangladesh, and South Africa, respectively (Figure 2A-E). The mean read counts of BAV were significantly higher in the samples from Bangladesh and South Africa compared to those from the UK (p < 0.001) (Figure 2E); however, its prevalence in term and preterm placentas was not significantly different. Our further examination of the sequence alignment on the human genome revealed a sequence homology of approximately 300 nt between BAV and a segment of the human genome; however, the origin of this homology and chromosomal integration remains unclear. Therefore, we disregarded BAV as a viral species for our subsequent analyses.

**Figure 2:**
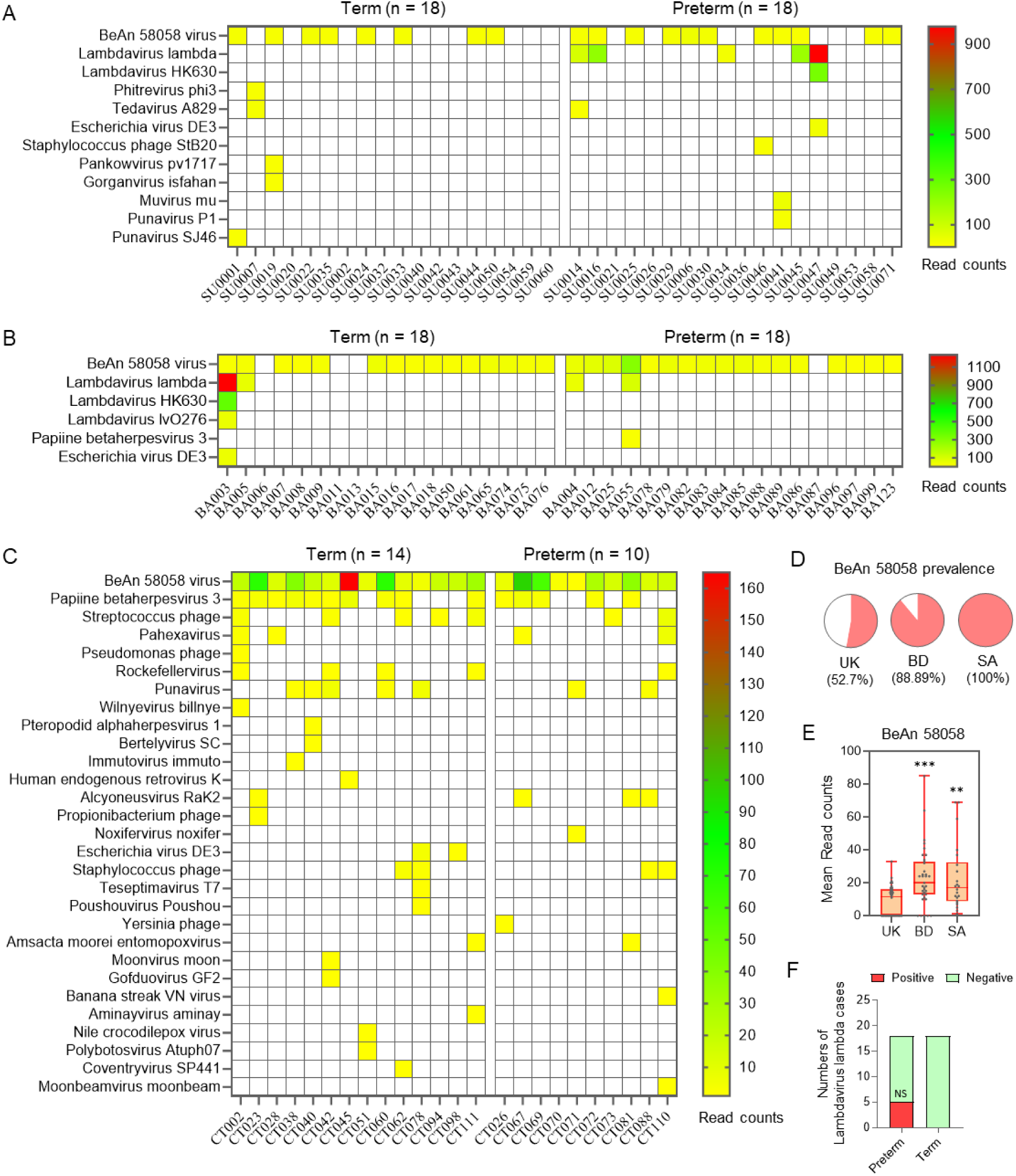
Placental virome profiles of women from the UK, Bangladesh, and South Africa in term and preterm births. Heatmaps displaying virus read counts in individual placentas from the UK (A), Bangladesh (B), and South Africa (C) cohorts. Blank squares signify no virus detection. D, E. Prevalence (D) and mean read counts (E) of BeAn 58058 viral transcripts in the UK (n = 36), Bangladesh (n = 36), and South Africa (n = 24) placentas. **p = 0.0044 UK vs. South Africa, ***p = 0.0008 UK vs. Bangladesh; One-way ANOVA with Tukey’s post hoc test. F. Prevalence of *Lambdavirus lambda* in term and preterm placentas within the UK cohort. NS = not significant, Fisher’s exact contingency test.

We identified *Lambdavirus lambda,* a dsDNA virus in the class *Caudoviricetes* (Escherichia phage lambda), in five UK (14%) and four Bangladeshi (11%) placenta samples, whereas it was not detected in any South African samples (Figure 2A, B, C). This virus was only detected in preterm placentas (27.78%) within the UK cohort, with an odds ratio of 8.14 (95% CI: 1.00-97.55, p = 0.09; Fisher’s exact test) (Figure 2A, F). In contrast, *Papiine betaherpesvirus 3* (PaHV-3), a dsDNA virus in the genus *Cytomegalovirus*, was detected in 62.5% of South African placentas, whereas only one case was found in the preterm group of Bangladeshi samples, with no cases in the UK cohort (Figure 2A, B, C).

Additionally, Streptococcus phage (29.1%), *Punavirus* (29.1%), and Staphylococcus phage (16.67%) were relatively more prevalent in the South African samples (Figure 2C); however, Fisher’s exact contingency test revealed no association between these viruses and PTB (data not shown). Other viruses were detected in the samples at low frequencies (Figure 2C).

Our virus detection sensitivity was optimised by identifying placenta-relevant DNA virus hCMV (human cytomegalovirus) and RNA virus ZIKV (Zika virus) in placenta samples spiked with these nucleic acids, employing single shotgun sequencing within the same sample. Our shotgun DNA sequencing effectively detected both hCMV and ZIKV in the spiked samples in a concentration-dependent manner (Figure S1A, B).

To investigate whether the placental viromes were contaminated by the vaginal microbiome during vaginal deliveries, we conducted shotgun metagenomic sequencing on a subset of placenta-linked cervicovaginal fluid (CVF) samples from eight women (n = 8) in the Bangladeshi cohort who delivered vaginally. The CVF samples exhibited relatively rich virome profiles compared to the placentas, with the most prevalent viral species being Lactobacillus phages (62.5%), *Gelderlandvirus melville* (62.5%), *Elpedvirus* (50%), *Triavirus* (a Staphylococcus phage) (37.5%), and Streptococcus phage (25%) (Figure 3A). The mean read counts for Staphylococcus phage *Triavirus* and Lactobacillus phages were significantly higher than those of other species (p < 0.05) (Figure 3B). None of the CVF virus species were detected in the linked placenta samples, suggesting that the detected placental viral species were less likely to result from contamination by the vaginal microbiota in the study samples.

**Figure 3:**
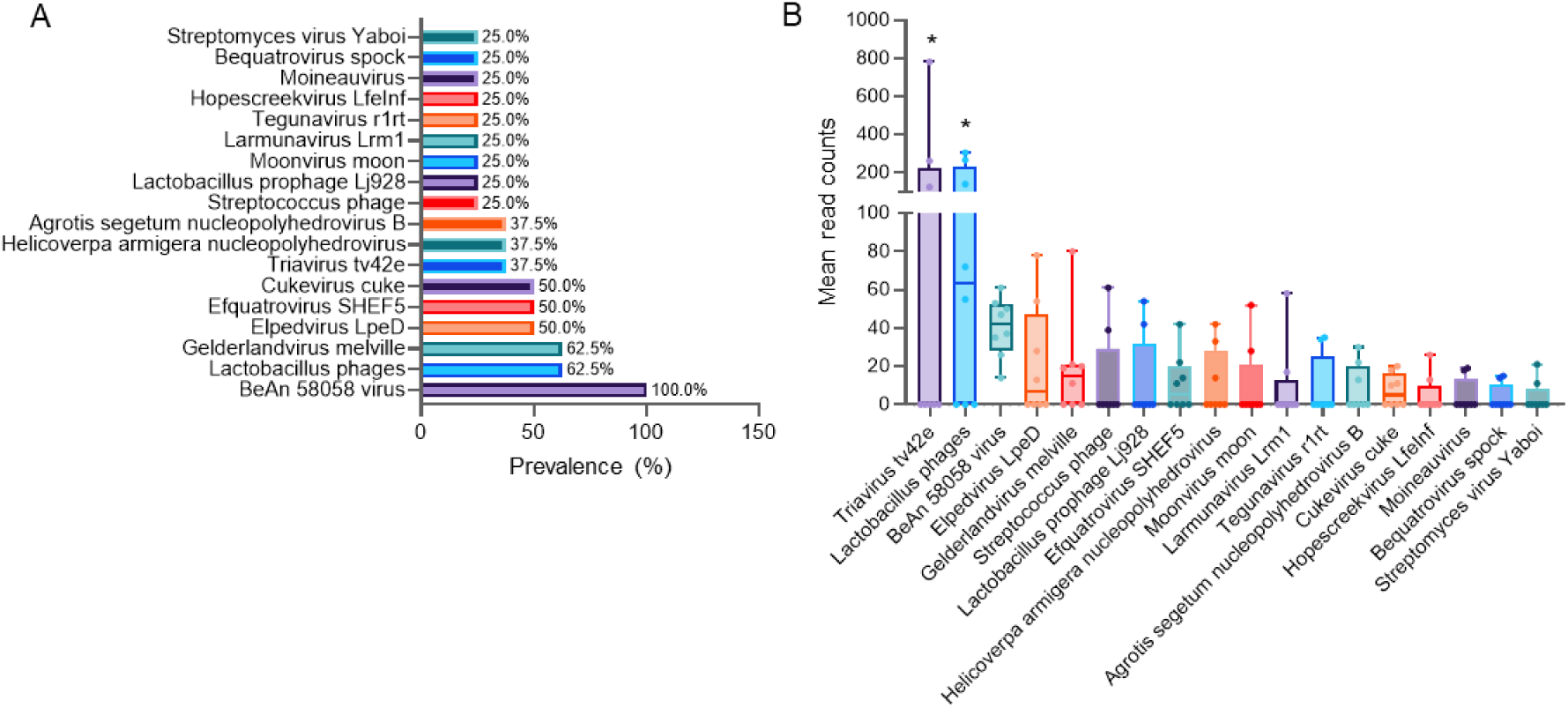
Cervicovaginal fluid (CVF) virome profile in eight Bangladeshi women. Bar diagrams showing prevalence (A) and mean read counts (B) of viruses in CVF samples (n = 8). *p < 0.05 vs other species. One-way ANOVA with Tukey’s post hoc test (B).

Due to the lack of linked CVF samples from the same subjects in the UK and South African cohorts, we were unable to replicate this cross-verification approach in these groups. Nevertheless, our parallel sequencing of instrument swabs, laboratory air samples, and RNA and DNA extraction buffers revealed no viruses, thereby ruling out the possibility of environmental or reagent viral contamination of the placental samples during processing. BAV was detected in all eight CVF samples, as observed in the placentas; however, it was not found in the buffer or laboratory environment controls, suggesting that the source of BAV is human host DNA.

In addition to shotgun sequencing, we conducted real-time qPCR array analysis targeting eight pathogens, including HSV-1, HSV-2, hCMV, HHV-6, HHV-7, Human Parvovirus B19, *Enterovirus,* and *Treponema pallidum,* on 175 Bangladeshi placenta tissue samples (119 term, 56 preterm), which included the samples used for shotgun sequencing (Table S1). The PCR analysis identified Human Herpesvirus 6 (HHV-6) in only one preterm placenta, with a Ct value of 25.25, and was therefore regarded as a positive detection. Human Betaherpesvirus 7 (HHV-7) was found in three term placentas, whilst Human Parvovirus B19 was detected in one term placenta, with intermediate Ct values of > 37.5 and 36, respectively, and were thus considered inconclusive detections (Table S1).

Overall, our data reveals a distinct pattern of placental viromes across the cohorts, showing low prevalence in the UK (27.78%) and Bangladesh (10.1%), but a higher prevalence among South African women (95.8%). Notably, *Lambdavirus lambda* was exclusively identified in preterm placentas from the UK, while the PaHV-3 species was highly prevalent among South African women, without any significant association with PTB. Various bacteriophages were sporadically detected in placentas across the UK, South Africa, and Bangladesh; however, the presence of pathogenic viruses in idiopathic spontaneous PTB was rare.

### 2.3. Diverse bacteriome profiles are present in the UK, South African and Bangladeshi placentas

Our primary aim in this study was not to profile the placental bacteriome; consequently, samples were treated with lysozyme/DNase I to eliminate non-enveloped nucleic acids, including host DNA, while preserving enveloped viral nucleic acids^25, 26^. Despite this initial treatment, our shotgun metagenomic sequencing identified a variety of bacterial species in 77.78% of UK, 58.33% of Bangladeshi, and 87.5% of South African villous/decidua placenta samples (Figure 4). The most prevalent species were *Moraxella osloensis*, *Delftia lacustris,* and *Cutibacterium acnes*, which appeared in varying abundances across the cohorts. For instance, *M. osloensis* (a gram-negative, aerobic bacterium) was found in 66.7% of UK and 50% of South African placenta samples, with mean abundance significantly higher in UK placentas compared to those from South Africa (718 reads/sample vs 50 reads/sample, p < 0.00001). Conversely, this species was not detected in Bangladeshi samples (Figure 4A, D, G). *D. lacustris* (a rod-shaped, gram-negative bacterium) was the dominant species (90.5% positive cases) in Bangladeshi placenta, where its mean abundance was significantly greater in preterm placentas compared to term placentas (p = 0.028) (Figure 4D, E, F). Of the 24 South African placentas, bacteria were identified in 21 samples (87.5%), all of which tested positive for *C. acnes* (a rod-shaped, gram-positive bacterium) with no specific association with preterm birth (Figure 4G). We also detected *Escherichia coli* (a gram-negative, rod-shaped bacterium) in four UK, four South African, and one Bangladeshi placenta. *E. coli* was exclusive to the UK’s preterm placentas, where its phage *Lambdavirus lambda* was identified within the same samples, demonstrating a significant positive correlation with *E. coli* (Spearman R = 0.99, p = 0.017) (Figure 4H). *Staphylococcus epidermidis* (gram-positive bacteria) was identified in two preterm placentas (5.6%) in the UK cohort and six placentas (25%) in the South African cohort, but none in the Bangladeshi cohort.

**Figure 4:**
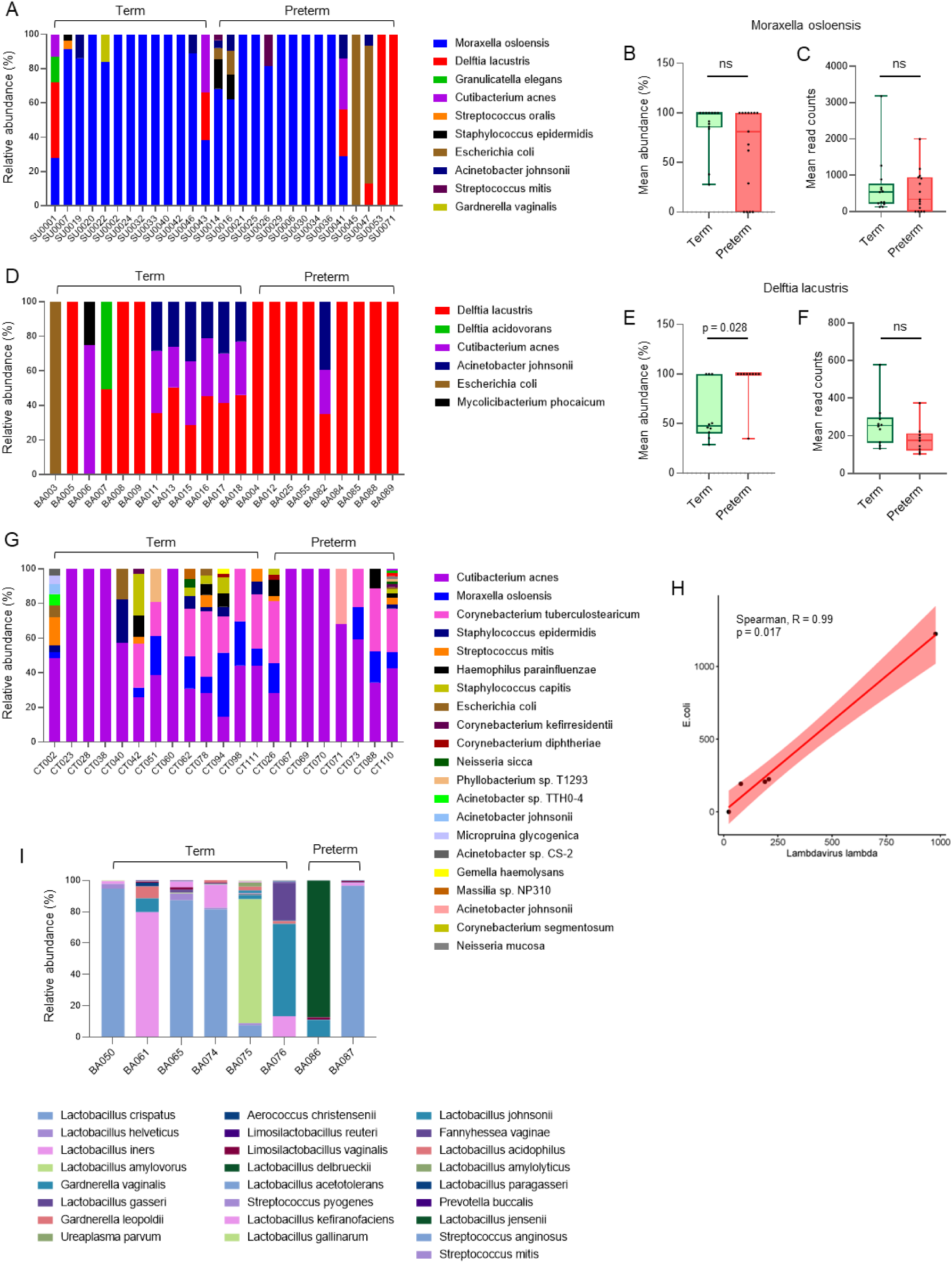
Bacterial species identified in placenta and cervicovaginal fluid (CVF) samples via shotgun metagenomic sequencing. A. Bar diagrams depicting the relative abundance of bacteria in UK placenta samples, along with mean abundance (B) and mean read counts (C) of *M. osloensis* in the term and preterm groups within the UK cohort. D. Relative abundance of bacteria in Bangladeshi placenta samples, accompanied by mean abundance (E) and mean read counts (F) of *D. lacustris* in the term and preterm placentas of the Bangladeshi cohort. G. Relative abundance of bacteria in South African placenta samples. H. Spearman correlation analysis between *Lambdavirus lambda* and *E. coli* read counts in the UK placentas. I. Relative abundance of bacteria in the CVF samples from the Bangladeshi cohort. The Mann-Whitney U test was conducted between term and preterm groups (B, C, E, F). ns = not significant.

Furthermore, our shotgun metagenomic analysis of placenta-associated CVF samples from the Bangladeshi cohort (n = 8), as previously noted, primarily detected various species of lactobacilli, alongside *Gardnerella vaginalis* (Figure 4I), as anticipated. *D. lacustris* and *C. acnes*, which were predominantly observed in placentas, were absent in the CVF samples. Similarly, lactobacilli or other common vaginal species identified in the CVF samples were missing in the vaginally delivered Bangladeshi placentas from which the CVF was sourced. This observation diminishes the likelihood of contamination of the placenta samples with the vaginal microbiome during normal vaginal delivery in Bangladeshi cases.

We were unable to apply a similar technique for the UK and South African samples to eliminate the possibility of placental contamination with the vaginal microbiome due to lack of placenta-linked CVF samples. We noted some bacterial species in our technical negative controls (blank swabs and buffer) for CVF samples, however, the read counts for those bacteria were significantly smaller than that found in the placental tissue sample, suggesting a possible contamination of our blank control samples with the placental nucleic acid elements, and not *vice versa* (Table S2).

Next, we calculated odds ratios (OR) to estimate the likelihood of detecting viruses or bacteria (at least one species per sample) in placentas delivered vaginally compared to those delivered by caesarean section, as well as in cases of spontaneous rupture of membranes (SROM/PPROM) compared to their absence (Figure 5). Placental contamination with vaginal microbiota is generally higher in instances of vaginal delivery. However, our analysis did not indicate an increased likelihood of detecting viruses or bacteria in vaginal deliveries when compared to caesarean sections, aside from the Bangladeshi cohort. In the Bangladeshi samples, the positive detection rate for bacteria was significantly greater in vaginal delivery cases than in those delivered by caesarean sections (OR = 30, 95% CI: 3.13–287.06, p = 0.003) (Figure 5B). Although the odds of detecting bacteria in placenta samples were higher for vaginal deliveries, the bacterial species identified in the placenta did not correspond with the predominant bacterial species found in the vaginal microbiota (Figure 4D, I). This finding suggests that the bacteria detected in the placentas of the Bangladeshi cohort are unlikely to be contaminants derived from the vaginal microbiota. Furthermore, no relationship was observed between the presence of viruses or bacteria in placentas from any of the three countries and the status of membrane rupture (Figure 5C, D).

**Figure 5:**
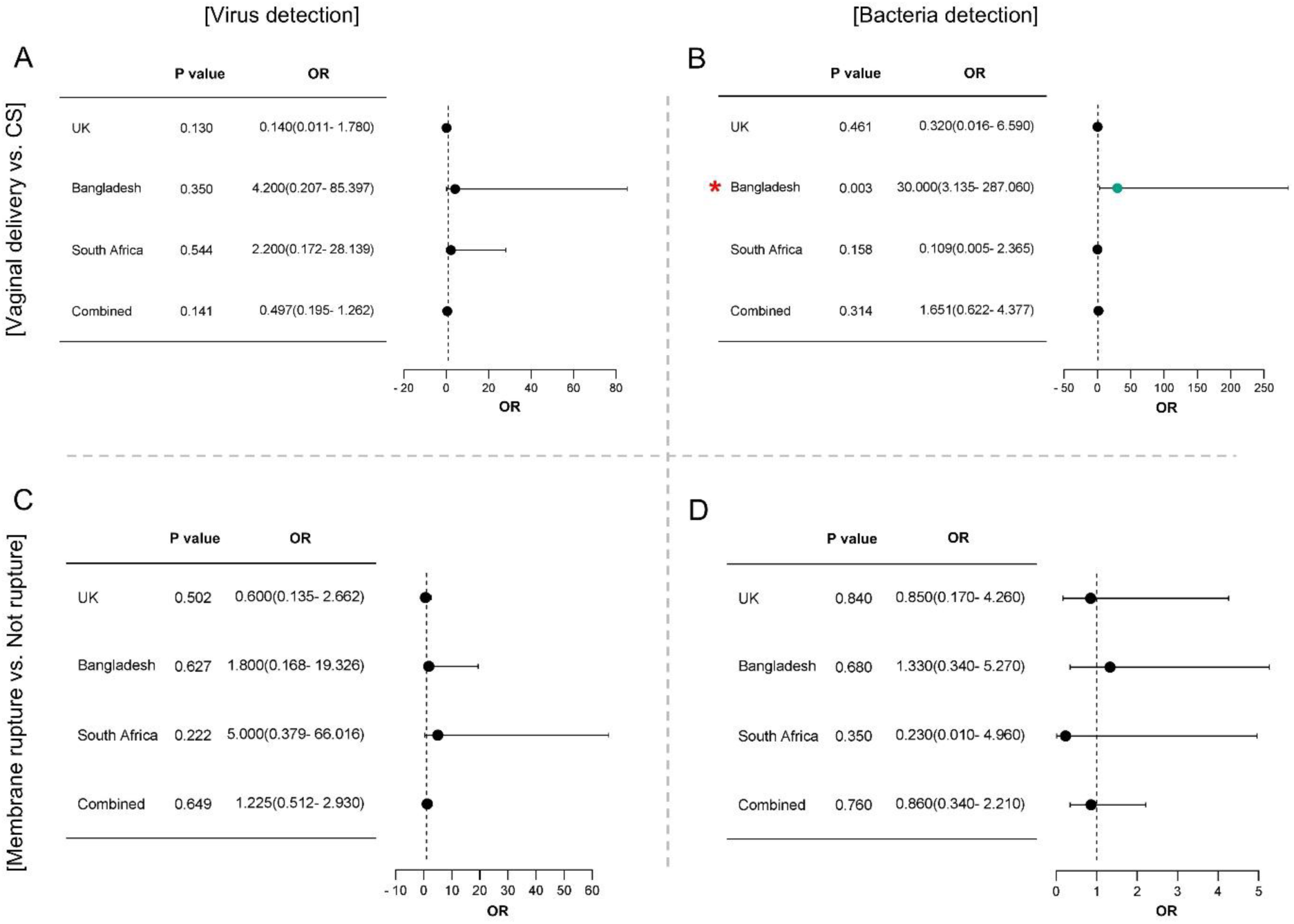
Forest plots illustrating odds ratios (OR) with 95% confidence interval (CI) ranges for detecting viruses (A, C) and bacteria (B, D) in vaginal deliveries (A, B) and in spontaneous rupture of membranes, regardless of term or preterm births. Detection of viruses and bacteria was conducted using shotgun metagenomic analysis. UK: n = 36, Bangladesh: n = 36, South Africa: n = 24.

Together, we conclude that the placenta exhibits diverse bacteriome profiles, characterised by the predominance of distinct non-pathogenic species among women from the UK, Bangladesh, and South Africa. However, many of these species may originate from the skin microbiota (e.g., *C. acnes*), although this could not be conclusively determined in our study.

### 2.4. Detection of intracellular bacteria within the villous placenta

To confirm whether the presence of bacteria in the placenta was due to contamination during labour, we fixed snap-frozen villous/decidua tissue blocks in 10% formalin. These blocks were excised from the same tissue biopsies used for shotgun metagenomic sequencing. The tissue blocks were embedded in paraffin, sectioned to a thickness of 5 µm, and stained using Gram and H&E staining methods. For this analysis, we selected five preterm placentas from the UK cohort that tested positive for *Lambdavirus lambda* and *E. coli* and compared them to five preterm placentas where *Lambdavirus lambda* and *E. coli* were not detected by shotgun sequencing. Additionally, we used five term placentas that tested negative for any bacteria by shotgun sequencing as a negative control.

Gram staining revealed intracellular gram-negative bacteria primarily within the chorionic villi of placentas where *E. coli* and *Lambdavirus lambda* were detected by shotgun sequencing (Figure 6A, 1st and 2nd columns). The gram-negative bacteria were localised within the cytoplasm of syncytiotrophoblasts, cytotrophoblasts, and inflammatory cells in the villous stroma (100x oil immersion, Figure 6A). In contrast, placentas in which *E. coli* was not detected by sequencing exhibited minimal or no Gram staining signals (Figure 6A, 3rd and 4th columns).

**Figure 6:**
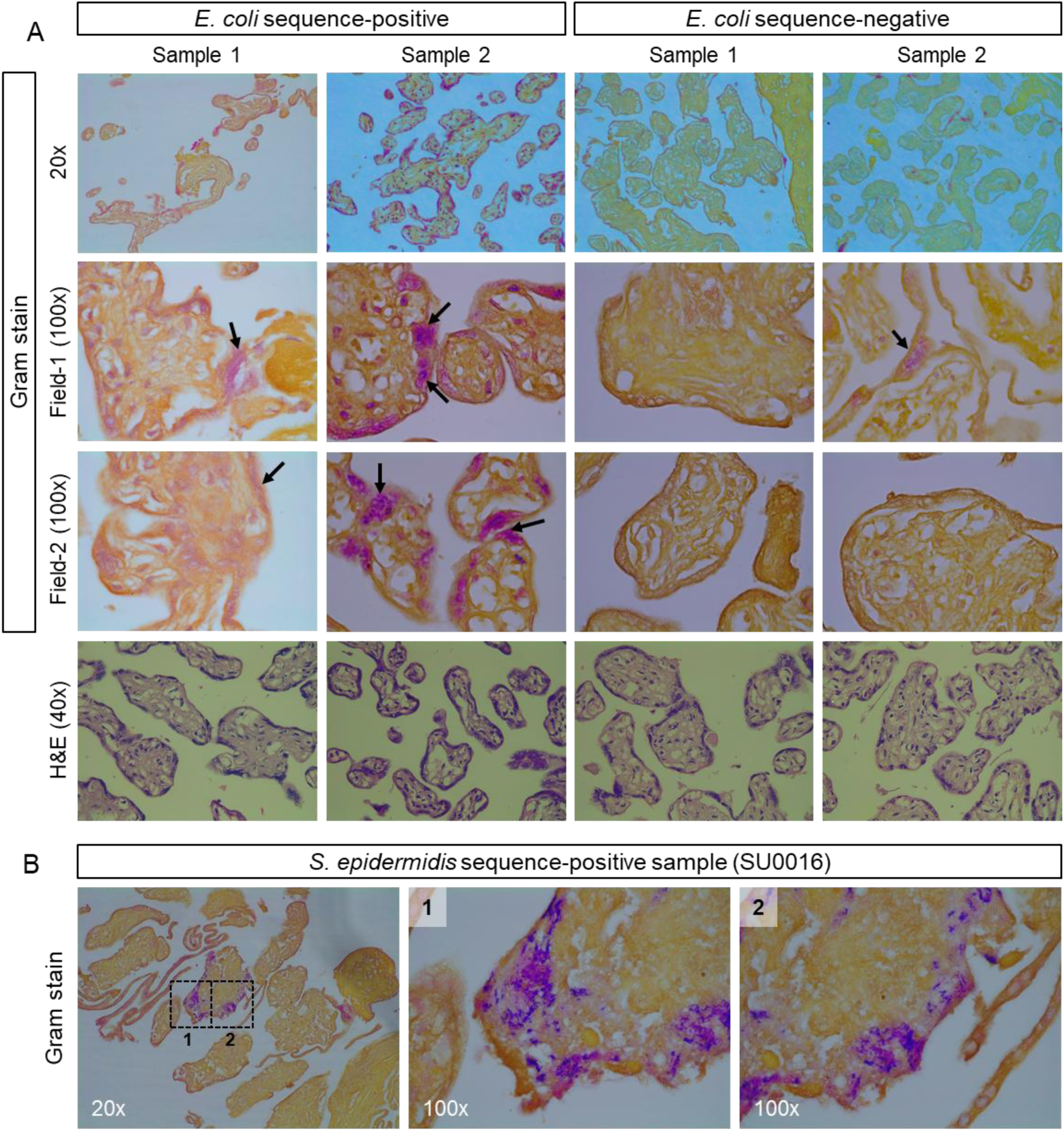
Gram and H&E staining of formalin-fixed paraffin-embedded placenta samples from the UK cohort. A. Representative microscopic images (20x and 100x oil immersion) from two independent samples showing positive staining (pink colour) for gram-negative bacteria in sequence-positive samples (1^st^ and 2^nd^ column), and low or negative staining for gram-negative bacteria in sequence-negative samples (3^rd^ and 4 column) (arrows). The bottom row shows H&E staining of the respective samples. B. Gram staining shows gram-positive (purple colour) and gram-negative (pink colour) within the villous tissue at 20x and 100x oil immersion images (1, 2).

Shotgun sequencing also revealed the presence of the gram-positive bacterium *Staphylococcus epidermidis* alongside gram-negative bacteria in two preterm placenta samples (SU0014 and SU0016) (Figure 4A). Gram staining confirmed the existence of both gram-positive and gram-negative bacteria in one of these samples (SU0016) (Figure 6B). However, H&E staining did not show significant inflammatory infiltration in the chorionic villi, regardless of the presence or absence of bacteria in the placentas (bottom row, Figure 6A). No bacteria were detected in the sequencing-negative term placenta samples via gram staining (Figure S2).

Together, the strong positive correlation between *Lambdavirus lambda* phage and its host *E. coli* (Figure 4H), coupled with the presence of intracellular gram-negative bacteria in the same placenta samples, suggests that the detection of *E. coli* in villous placenta samples is unlikely to result from contamination originating from the lower gut or during the labour process. In addition, pathogenic bacteria, such as *S. epidermidis*, can occasionally be found in the UK preterm placentas.

### 2.5. *E. coli* and *Lambdavirus lambda* did not elicit significant immune response in placental villous tissue

To assess whether the presence of gram-negative bacteria (e.g., *E. coli*) and its phage *Lambdavirus lambda* in the villous tissue of preterm placentas influences gene expression, we performed differential gene expression (DE) and gene set enrichment analysis (GSEA) on bulk RNA-seq data. These data were derived from identical villous placenta tissues identified as positive for both *E. coli* and *Lambdavirus lambda* (n = 4) and negative for both (n = 4) (GEO accession GSE211927; Table S3). The RNA-seq dataset was previously published in our earlier study^27^.

Given the small sample size and variations in library size, we conducted a weighted limma-voom differential gene expression analysis with robust settings to minimise biases from outliers^28, 29^. This analysis identified 34 significantly upregulated and 125 significantly downregulated genes (DEGs) in *E. coli*/*Lambdavirus lambda*-positive samples compared to *E. coli*/*Lambdavirus lambda*-negative samples (Log2FC > ±1.5, FDR < 0.05) (Figure 7A, B; Table S4). Notably, genes associated with extravillous trophoblast cells, cell cycle regulation, cell proliferation, and immune modulation (e.g., *HLA-G, NUF2, ICAM2, IL3RA*) were significantly downregulated in *E. coli*/*Lambdavirus lambda*-positive samples (Figure 7B; Table S4).

**Figure 7:**
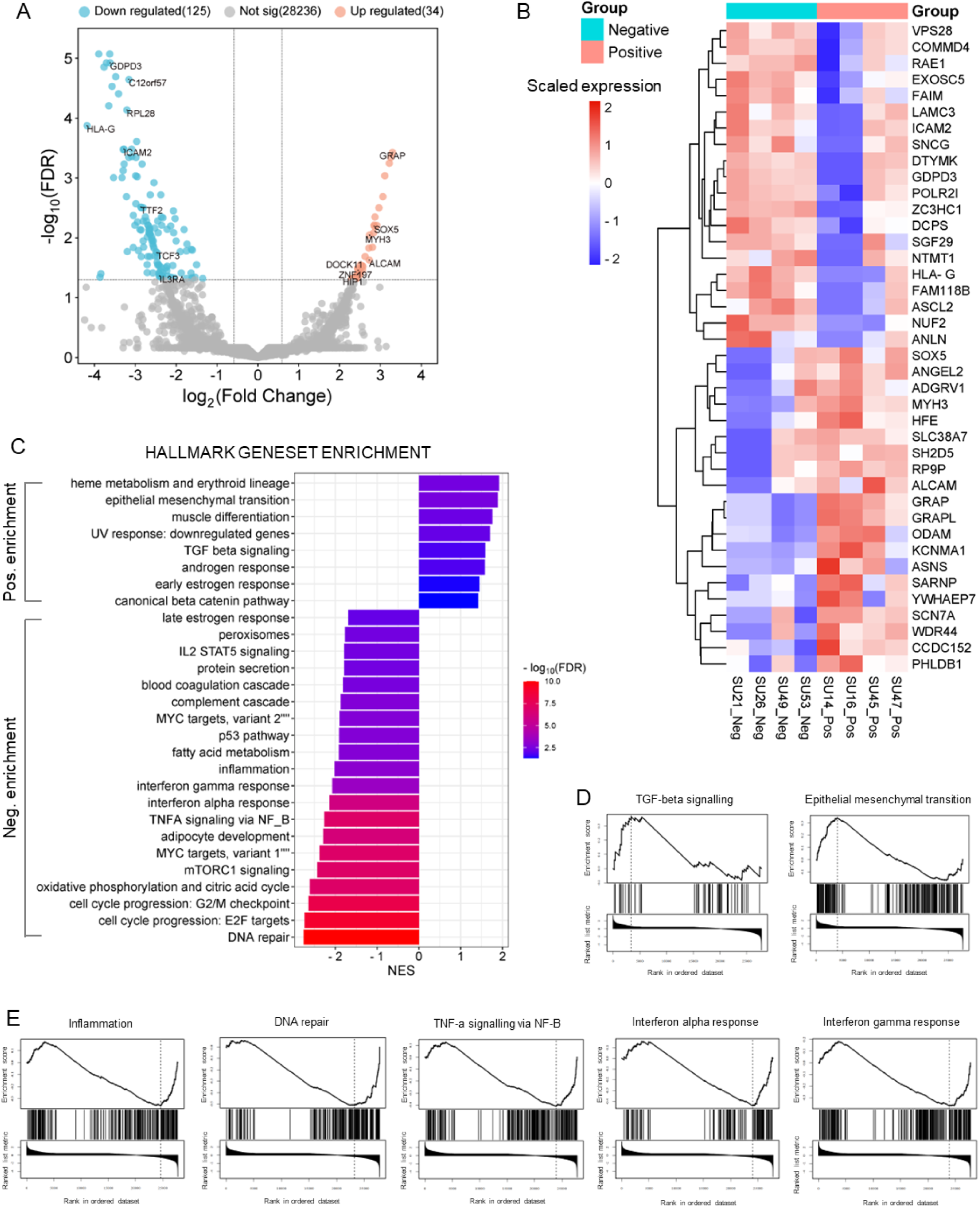
Bioinformatics analyses of RNA-seq data from the *Lambdavirus lambda/E.coli-*positive (n = 4) and - negative placentas (n = 4). A. Volcano plot showing upregulated and downregulated DEGs. B. Heatmap showing 20 upregulated and 20 downregulated DEGs with their expression in *Lambdavirus lambda/E.coli-*positive and - negative placenta samples. C. Bar diagram showing significantly positive and negative gene set enrichment of signalling pathways (FDR < 0.05) with normalised enrichment scores (NES) between *Lambdavirus lambda/E.coli-*positive and -negative placenta by GSEA. D, E. Enrichment score strips from selected positive enrichment (D) and negative enrichment (E) of gene sets.

GSEA was conducted using the Hallmark gene set database from the Human MSigDB Collections^30^. The analysis revealed a negative enrichment of signalling pathways typically associated with bacterial and viral infections, including inflammation, TNF-alpha signalling, interferon-alpha response, and interferon-gamma response (Figure 7C, E). Conversely, upregulated pathways included epithelial-mesenchymal transition (EMT), TGF-beta signalling, early oestrogen response, androgen response, and the canonical beta-catenin pathway.

This gene set enrichment pattern suggests that the presence of bacteria and bacterial phages within placental tissue did not provoke significant immune responses but could be associated with trophoblast dysfunction driven by EMT, TGF-β and Wnt signallings. Based on these findings, we conclude that *E. coli*, *S. epidermidis*, and *Lambdavirus lambda* in placental tissues are more likely to be non-immunogenic.

## 3. DISCUSSION

Most previous microbiome studies on human placentas have primarily utilised genomic DNA or 16S rRNA sequencing. These methods are restricted to detecting bacteria and DNA viruses. In this study, we employed shotgun metagenomic sequencing by separately extracting DNA and RNA from the same placenta tissues. The extracted RNA was reverse-transcribed into dsDNA and combined with genomic DNA prior to library preparation for nanopore next-generation sequencing. This methodology allowed for the simultaneous detection of DNA and RNA viruses, as well as bacterial species, in a single sequencing run. In the validation assay, we demonstrated that this approach can detect both DNA virus (eg. hCMV) and RNA virus (eg. ZIKV) in the same sample in a single sequencing in a dose-dependent manner.

Using this novel technique, we did not detect any known pathogenic viruses in any of the 96 placenta samples collected from uncomplicated term and preterm births in cohorts from the UK, Bangladesh, and South Africa. This sequencing result was corroborated by our real-time PCR array analysis of 175 Bangladeshi placenta samples, which similarly showed no positive detection of a selected panel of eight pathogenic organisms, including seven viruses, apart from one preterm placenta where HHV-6 was detected. HHV-6 has been implicated in placental infections and adverse pregnancy outcomes, including preterm birth. The virus can infect trophoblast cells, triggering inflammatory responses and immune modulation that disrupt placental function^31, 32^.

In our study, the preterm baby born alive with HHV-6 placental infection, without any maternal clinical manifestations, had a birth weight of 2.4 kg, which was slightly higher than the average birth weight of preterm babies in the Bangladeshi cohort (2.27 kg). The gestational age (GA) was also marginally higher (35.7 weeks) compared to the average GA for our PTB samples (34.6 weeks). This finding suggests that HHV-6 may persist in a dormant state within the placenta in less than 1% of Bangladeshi women, with no significant clinical impact at birth. Our data indicate that placental infections with known pathogenic viruses in uncomplicated term and preterm births are rare occurrences.

The placenta may harbour various viruses, primarily bacteriophages, which are characterised by low abundance and low prevalence. We observed a preferential dominance of certain viral species across the three countries studied. For instance, the *E. coli* phage *Lambdavirus lambda* was detected in placenta samples from the UK and Bangladesh but not from South Africa, despite its host, *E. coli*, being found in four South African placentas. One plausible explanation for this discrepancy could be the significantly lower abundance (28-fold fewer read counts) of *E. coli* in South African samples compared to those from the UK. However, this hypothesis does not apply to the Bangladeshi samples, as *Lambdavirus lambda* was observed in three samples where *E. coli* was not present. In contrast, we noted a linear relationship between the presence of *Lambdavirus lambda* and *E. coli* in the UK samples, supporting the established notion that *E. coli* is the primary host of *Lambdavirus lambda*^33, 34^. The independent presence of *Lambdavirus lambda* in the absence of its natural host in Bangladeshi placentas could be due to complete destruction of *E. coli* DNA by lysozyme/DNase I treatment during VLP DNA purification step.

Additionally, we observed the exclusive presence of both *Lambdavirus lambda* and *E. coli* in preterm placentas, but not in term placentas, within the UK samples. Although *Lambdavirus lambda* does not directly interact with human tissue, it can integrate into the chromosome of its host, *E. coli*, through the lysogenic cycle, potentially enhancing bacterial virulence^35^. However, our bioinformatics analysis did not identify any notable inflammatory or immune responses in tissue samples containing *Lambdavirus lambda*/*E. coli* compared to those where they were absent. This observation suggests that *E. coli* resides in the placenta as a normal commensal. The presence of *E. coli* as a normal commensal in the human placenta has been previously reported^4, 6^.

However, an earlier study suggested that the detected *E. coli* might have originated from contamination, as it was present in the majority of samples at high abundance, and all genomic signatures matched a particular strain^6^. In our study, *E. coli* was found in only specific samples, despite all being sequenced together. It is less likely that the *E. coli* originated from the endometrium or vaginal lining, as it has not been identified as a normal commensal in these anatomical regions but could be present occasionally^10, 36^. Furthermore, gram staining revealed the presence of gram-negative bacteria deep within the placental tissue, ruling out the possibility of contamination during labour.

Another viral species, *Papiine betaherpesvirus 3* (PaHV-3), also known as *Cytomegalovirus papiinebeta-3*, which is equivalent to *Baboon cytomegalovirus OCOM4-37*^37^, was detected in over 60% of South African placentas, one Bangladeshi placenta, and none of the UK samples. PaHV-3 exhibits genetic similarities to hCMV and HHV-6 and shares critical characteristics, including latency, reactivation and a predilection for placental tissue. While direct evidence of PaHV-3 infection and its impact on the human placenta remains limited, studies in primates have demonstrated its ability to infiltrate reproductive tissues and establish latency^24, 38, 39^.

In our study, PaHV-3 was detected in both term and preterm placentas, with no statistically significant association with PTB. The origin of PaHV-3 in South African placentas remains unclear; however, it is unlikely that the virus originated from reagents, instruments, or laboratory environmental sources, as control samples tested negative for PaHV-3. Furthermore, we found no association between the presence of any viral species in placenta samples and delivery mode or membrane rupture status in any of the three cohorts studied.

While our primary aim was to define virome profiles in the placenta, we also detected several bacterial species across the UK, Bangladeshi, and South African samples. In addition to *E. coli*, we identified the gram-positive bacterium *Staphylococcus epidermidis* in two preterm placentas from the UK cohort and six placentas from the South African cohort. *S. epidermidis* has previously been reported in human placentas as a normal commensal^4^. Interestingly, *Moraxella osloensis* was commonly detected in placentas from the UK and South African cohorts but was absent in the Bangladeshi samples. *M. osloensis* is a gram-negative opportunistic human pathogen and is saprophytic on the skin and mucosa. While it rarely causes human diseases, it has been associated with a range of conditions, including meningitis, vaginitis, sinusitis, bacteraemia, endocarditis, peritonitis, ocular infections and septic arthritis^40, 41, 42, 43, 44, 45, 46, 47, 48, 49, 50^.

In our study, *M. osloensis* was detected across both term and preterm placentas, with no significant association with PTB. However, its abundance was 14-fold higher in the UK samples compared to the South African samples. While *M. osloensis* is part of the skin microbiota^48^, it has not been identified as a component of the microbiota in the female reproductive tract^10^. This suggests that *M. osloensis* is unlikely to have originated from the uterus. Furthermore, the high abundance of this bacterium in the UK placentas may not be solely attributable to contamination from maternal skin. There is, however, a possibility of contamination from hospital environments (e.g., labour rooms and operating theatres) in the UK, as *M. osloensis* has been identified as a common airborne pathogen in indoor environments^51^. Notably, environmental air swabs from our laboratory, where placentas were processed, tested negative for any viruses or bacteria. Further studies are required to better understand the origin of *M. osloensis* in the placenta and its potential contributions to adverse pregnancy outcomes, as this bacterium can occasionally cause severe diseases.

In contrast, *Delftia lacustris* was detected in the majority of Bangladeshi placentas at high abundance and in a few UK placentas, but it was absent in South African placentas. *D. lacustris* is a gram-negative, aerobic, motile bacillus, which have been associated with human disease^52, 53, 54, 55^. In our study, the abundance of *D. lacustris* was significantly higher in preterm placentas compared to term placentas, although its potential contribution to the mechanisms underlying PTB remains unexplored. Notably, *D. lacustris* has recently been isolated from a vaginal discharge sample in a patient with cervical intraepithelial neoplasia^56^, and other *Delftia* spp. have been reported as part of the microbiota in the endometrium and fallopian tubes^10^.

Additionally, we observed a significant relationship between vaginal delivery and the presence of bacterial species in placentas within the Bangladeshi cohort, though the associated uncertainty was high. To investigate the potential origin of *D. lacustris* in placentas, we sequenced vaginal fluid samples from eight patients who delivered vaginally and whose placentas were subjected to shotgun sequencing. *D. lacustris* and other bacteria or viruses detected in the placenta samples were absent in these vaginal fluid samples. This finding suggests that the vagina was an unlikely source of *D. lacustris*, instead we suspect that the bacteria may arise from the endometrium.

Finally, *Cutibacterium acnes* was detected in placenta samples from all three countries, with its abundance and prevalence being notably higher in the South African samples. *C. acnes* is part of the microbiota of the skin, oral cavity, large intestine, conjunctiva, and external ear canal^57^. Thus, its presence in the placenta may result from contamination during labour. However, the markedly lower prevalence of this bacterium in UK samples compared to South African samples is intriguing. Although typically a commensal organism, *C. acnes* can act as an opportunistic pathogen^58^. In our study, we found no significant association between *C. acnes* and PTB.

While our technique identified several bacterial species in the placenta, some of these detections may be attributable to the transfer of bacterial extracellular vesicles (EVs) containing bacterial nucleic acids from maternal microbiota communities in other organs, such as the gut, to the placenta, which albeit requires further investigation^59, 60^.

Microbiome profiling of the placenta presents unique challenges owing to its low biomass and the risk of contamination during labour^8, 9^. To tackle these issues, we established several checkpoints in our study. Initially, we processed two batches of control samples alongside placenta and CVF sequencing. In one batch, we collected swabs from instruments, laboratory air, and DNA and RNA extraction reagents. Identical buffers were employed for processing samples from all three countries. These four control samples were sequenced concurrently with the placenta samples. No viruses or bacteria were identified in these four control samples. However, on a separate occasion when CVF samples were sequenced, we observed some organisms in the buffer control. The abundance of these bacterial species and viruses was significantly lower in the buffer control compared to the CVF samples, indicating a possible contamination of the control with sample material.

Our second strategy was to establish a relationship between the mode of delivery and the detection of viruses and bacteria in placenta samples. This approach suggested a reduced likelihood of placental contamination by vaginal microbiota. As a third strategy, we cross-compared our data across the three countries. Despite employing a standardised protocol for tissue and nucleic acid processing and sequencing, we observed distinct signatures of the virome and bacteriome with varying prevalence across samples from the UK, Bangladesh, and South Africa.

**Together, we conclude:** (i) The human placenta harbours a distinct virome with low prevalence and low abundance, primarily composed of bacteriophages across three ethnic backgrounds from three geographical regions. Bacteriophages such as *Lambdavirus lambda* is more prevalent in UK and Bangladeshi placentas, while *Punavirus* is more common in South African placentas. The PaHV-3 virus is highly prevalent in South African placentas without any significant association with PTB. (ii) The presence of pathogenic viruses in the placenta is rare and may not be associated with the pathogenesis of uncomplicated spontaneous PTB. (iii) The placenta contains a diverse array of bacterial species, including *M. osloensis*, *D. lacustris*, and *C. acnes*, with higher prevalence in the UK, Bangladeshi, and South African placentas, respectively. *E. coli* and *S. epidermidis* are also present in placenta samples, albeit at low frequencies, with *E. coli* being exclusively detected in UK preterm placentas. Further studies are needed, particularly focusing on *M. osloensis* and *D. lacustris*, to better understand their origin in the placenta and their potential association with adverse pregnancy outcomes.

## 4. METHODOLOGY

### 4.1. Placenta tissue harvesting

Placentas were collected in either a clean bucket or a sterile Ziplock plastic bag, and tissue samples were processed within 3 hours of delivery. Tissue harvesting was performed under sterile conditions in a biosafety level 2 (BSL-2) laminar flow hood. Sterile disposable draping and dissecting tools were used to minimise the risk of contamination. Two full-thickness tissue biopsies (1 cm × 1 cm × 2 cm) were obtained from the healthy-appearing, intact pericentral lobe of the placenta, which includes the decidua basalis and chorionic villous tissue (Figure S3). The biopsies were placed in sterile cryovials and immediately snap-frozen in liquid nitrogen for 1 hour before being transferred to a -80 °C freezer for later use. One biopsy was allocated for DNA extraction, and the other was designated for RNA extraction. A flowchart of tissue harvesting to subsequent metagenomic sequencing and data analysis is presented in Figure S3.

### 4.2. Viral-like particles (VLPs) enrichment and DNA extraction from the placenta

VLP enrichment and DNA extraction from the human placenta were conducted following previously published protocols with minor modifications^25, 26, 61^. Briefly, snap-frozen biopsies were thawed, and 50 mg of tissue was excised from the decidua basalis and 50 mg from the villous tissue. These tissue samples (totalling 100 mg) were minced on a sterile Petri dish under sterile conditions. The minced tissue was transferred to a sample shredder tube (Qiagen, Cat No. 990381) containing a 5 mm stainless steel bead (Qiagen, Cat No. 69989) and 1 ml of sterile digestion buffer (1 mg/ml collagenase D + 1 U DNase I in PBS).

The samples were incubated at 37 °C for 1 hour, with intermittent shredding for 2 minutes at 50 Hz using a TissueLyser LT every 20 minutes. Following incubation, the tissue was further shredded at the same speed for 5–10 minutes to ensure complete lysis. The lysed samples were centrifuged at 5,000 × *g* for 5 minutes at room temperature, and the supernatants were transferred to new 1.5 ml sterile Eppendorf tubes.

Supernatants were treated with lysozyme (1 mg/ml) (ThermoFisher, Cat No. 90082) at 37 °C for 30 minutes to lyse bacterial and host cell membranes. Subsequently, 0.2x volume of chloroform was added to each sample, vortexed for 1 minute, and incubated at room temperature for 10 minutes to degrade residual bacterial and host cell membranes, exposing the nucleic acids. The samples were then treated with DNase I (ThermoFisher, Cat No. AM2238) (10 U per ml of sample) at 37 °C for 10 minutes, followed by heat inactivation at 65 °C for 10 minutes. This process effectively degraded most host and bacterial DNA without significantly impacting the DNA within the VLPs^25, 26^.

To extract viral DNA, VLP-containing samples were divided into two Eppendorf tubes (400 µl per tube) and lysed using 4% SDS (ThermoFisher, Cat No. 15553027) and 40 mg/ml Proteinase K (Qiagen, Cat No. 19131) at 56 °C for 20 minutes. Following lysis, 60 µl of 5 M NaCl (ThermoFisher, Cat No. AM9760G) and 70 µl of 10% (wt/vol) cetyltrimethylammonium bromide (CTAB)/0.7 M NaCl solution (Merck, Cat No. H6269-100G) were added to each tube, vortexed for 30 seconds, and incubated at 65 °C for 10 minutes. The samples were then mixed with an equal volume (750 µl) of ultra-pure phenol:chloroform:isoamyl alcohol (25:24:1) (ThermoFisher, Cat No. 15593031), vortexed for 30 seconds, and centrifuged at 16,000 × *g* for 5 minutes at room temperature. The aqueous phase (∼450 µl per tube) was collected into a sterile 15 ml centrifuge tube, yielding ∼900 µl per sample. For ethanol precipitation, each sample was supplemented with 1 µl of glycogen (20 µg/ml) (ThermoFisher, Cat No. 10814010), 0.5x volume of 5 M NH_4_OAc (ThermoFisher, Cat No. AM9070G), and 2.5x volume of 100% ethanol. The mixture was inverted to mix and incubated at -80 °C for at least 1 hour.

Samples were then centrifuged at 16,000 × *g* for 30 minutes at 4 °C to pellet the DNA. The supernatants were removed, and 150 µl of 70% ethanol was added to each sample. After gentle mixing, the samples were centrifuged at 16,000 × *g* for 2 minutes at 4 °C. The ethanol was carefully removed, and the DNA pellets were air-dried before being resuspended in 100 µl of RNase/DNase-free water. The extracted DNA was subsequently cleaned and concentrated using the DNA Clean & Concentrator-5 kit (Zymo Research, Cat No. D4013) in accordance with the manufacturer’s protocol and eluted in 10 µl of DNA Elution Buffer. The final DNA concentrations ranged from 65–100 ng/µl, as quantified using the Qubit dsDNA HS Assay Kit (Invitrogen, Cat No. Q32851) on a Qubit fluorometer. Samples were stored at -80 °C for subsequent processing.

### 4.3. RNA extraction from placenta

Total RNA was extracted from snap-frozen placentas using the RNeasy Plus Mini Kit (Qiagen, Cat. No. 74134) following the manufacturer’s instructions and a previously published protocol^27, 62^. Briefly, frozen tissue biopsies designated for RNA extraction were thawed, and 40 mg of tissue was excised from the villous and decidua basalis regions (in a 50:50 ratio). The tissue samples were placed into a sample shredder tube containing a 5 mm stainless steel bead and RLT Lysis Buffer supplemented with β-mercaptoethanol (Sigma, Cat No. M3148).

Tissue homogenisation was performed using a TissueLyser LT homogenizer for 10 minutes at 50 Hz. The resulting homogenate was centrifuged at 18,000 x *g* for 3 minutes, and the supernatant was passed through a gDNA eliminator spin column to remove genomic DNA. Total RNA was extracted using a mini spin column according to the manufacturer’s protocol. The purified RNA was eluted in 50 µl of RNase/DNase-free water. The RNA yield was quantified using the Qubit RNA HS Assay Kit (ThermoFisher, Cat No. Q32852). RNA samples were stored at -80 °C for subsequent use.

### 4.4. dsDNA synthesis and purification

Double-stranded DNA (dsDNA) was synthesised from total RNA using the Maxima™ H Minus Double-Stranded cDNA Synthesis Kit according to the manufacturer’s instructions (ThermoFisher, Cat No. K2562). Briefly, for first-strand cDNA synthesis, 5 µg of total RNA was combined with 1 µl of random hexamer primers and incubated at 65 °C for 5 minutes, followed by immediate chilling on ice. Subsequently, 5 µl of 4x First Strand Reaction Mix and 1 µl of First Strand Enzyme Mix were added to each reaction. The reaction mixture was incubated at 25 °C for 10 minutes, followed by 50 °C for 30 minutes, and terminated by heating at 85 °C for 5 minutes before being placed on ice.

For second-strand cDNA synthesis, 20 µl of the first-strand cDNA reaction mix was used in a 100 µl reaction volume. The mixture was incubated at 16 °C for 1 hour following the manufacturer’s protocol (ThermoFisher, Cat No. K2562). The reaction was terminated by adding 6 µl of 0.5 M EDTA, and residual RNA was removed by treating the sample with 10 µl of RNase I (100 U) for 5 minutes at room temperature. Following the manufacturer’s protocol, the resulting dsDNA was cleaned and concentrated in 25 µl of Elution Buffer using the GeneJET PCR Purification Kit (Thermo Scientific, Cat No. K0701). The dsDNA samples were stored at -80 °C for future use.

### 4.5. Sample preparation for shotgun metagenomic sequencing

The entire yield of dsDNA (70-100 ng) was combined with 500 ng of VLP DNA, concentrated to 12 µl in Elution Buffer using the DNA Clean & Concentrator-5 kit as described above, and submitted for sequencing (Figure S3). For reagent controls during RNA and DNA extraction, sterile PBS was used in place of tissue and subjected to the same protocols for RNA and DNA extraction and dsDNA synthesis. For instrument and laboratory air controls, sterile Dacron swabs (Delta Lab, Cat No. 300263) were processed using the RNA extraction and dsDNA synthesis protocols. One control from each category was included in each batch of sample preparation.

For the viral RNA/DNA spiking assay, 5 µg of total RNA extracted from Bangladeshi placenta samples was spiked with genomic RNA from Zika virus (ZIKV) (ATCC, VR-1838DQ) at various copy numbers ranging from 5 x 10^5^ to 6.2 x 10^3^ copies per sample (Figure S1). These spiked RNA samples (n = 7) and one non-spiked RNA sample (0 copies/sample) were synthesised into dsDNA using the protocol described above. The resulting ZIKV-spiked samples were subsequently spiked with genomic DNA from Human herpesvirus 5 (human cytomegalovirus, hCMV) strain AD-169 (ATCC, VR-538DQ) at the same copy numbers per sample as ZIKV (Figure S1). These ZIKV/hCMV spiked dsDNA/DNA mixtures were then concentrated as above and submitted for sequencing.

### 4.6. DNA extraction from CVF samples

Bacterial DNA was extracted from eight CVF samples (n = 8) from Bangladeshi cohort along with a blank swab sample as negative control (to control for microbial contaminants in reagents) following our previously published protocol^63^. Prior to DNA extraction, all tips, tubes, tweezers and scissors were UV sterilised for 15 minutes to reduce bacterial DNA contamination. 500 µl sterile Phosphate buffered saline (PBS) was added to the thawed vaginal swab and vortexed for 5 minutes to elute the CVF for DNA extraction. The swab was placed into a new tube and centrifuged at 10,000 x *g* for 1 minute to draw out any remaining fluid from the swab. The 500 µl of CVF was transferred to a clean Eppendorf tube and incubated with 75 µl of 20 mg/mL lysozyme (Fisher Scientific, Cat No. 89833) at 37°C for 1 hour in order to degrade bacterial cell wall. DNA was extracted using the QIAmp DNA mini kit (Qiagen, Cat No. 51304) according to the manufacturer’s instructions adapting the protocol in our previous study^63, 64^. The purified DNA was eluted from the QIAamp kit spin column in 50 μl of buffer AE and quantified using the Qubit dsDNA HS Assay Kit (Invitrogen, Cat No. Q32851) on a Qubit fluorometer before storage at −80°C until further processing.

### 4.7. DNA sequencing

Sequencing libraries were prepared using 5 ng of template DNA and with Oxford Nanopore Technologies Rapid PCR Barcoding Kit (Cat No. SQK-RPB004) and a modified protocol to optimise results with viral RNA and DNA^65^. Sample libraries were individually barcoded, pooled into groups of 12 and sequenced with R9.4.1 flow cells (FLO-MIN106) on the GridION MK1 platform. Sequencing was run for 72 hours, with flow cells washed using the Flow Cell Wash Kit (Cat No. EXP-WSH004) and re-loaded with a fresh library aliquot after 36–48 hours to maximise yield. Basecalling was performed on-instrument using the high-accuracy model (MinKNOW v22.10.5; Guppy v6.3.8).

### 4.8. Real-time PCR array

For real-time PCR, 0.56 grams of tissue were treated with 1 mL lysis buffer (NucliSENS easyMAG Lysis Buffer, Cat. No. 280134, Biomerieux, France) and homogenized using 5 mm stainless steel beads (Cat. No. 69989, Qiagen, Germany). Nucleic acids were extracted using the QIAamp DNA Blood Mini Kit (Cat. No. 51106, Qiagen, Germany). Commercial real-time PCR kits were used for the detection of HSV-1, HSV-2, Treponema pallidum (FTD Genital Ulcer (REF. FTD-19-64-RUO, Fast-track Diagnostics, Luxembourg), Cytomegalovirus (FTD cytomegalovirus (REF. FTD-8.1-64-RUO, Fast-track Diagnostics, Luxembourg)), HHV-6, HHV-7, human parvovirus B19, and enterovirus (FTD Fever and Rash (REF. FTD-10.3-64, Fast-track Diagnostics, Luxembourg) using the C1000™ Thermal Cycler. PCR was performed according to the manufacturer’s instructions, and hRNP3 human RNase P was detected from each sample as an extraction control.

### 4.9. Bioinformatics analysis for pathogen detection

Sequencing was conducted in two separate batches for each sample (see Section 4.7). The raw *fast5* files were basecalled using ONT Guppy v6.3.8. The barcodes were filtered using Porechop v0.2.4 (--discard_middle)^66^. Quality-filtered reads were mapped to human genome (GRCh38.p13.genome.fa) using minimap2 v2.24 (-x map-out -a)^67^. Unmapped reads were retained for downstream analysis using samtools v1.16.1 and subsequently filtered with NanoFilt v2.8.0 (-q 10 -l 500 --headcrop 30)^66, 68^. Filtered reads from both batches of each sample were merged and taxonomically classified using the default settings of Kraken2 v2.1.2 against the ‘’Viral” and “Standard-8” databases independently^69^. Abundances of each taxon were calculated to species level using Bracken v2.8 (-l S)^70^.

### 4.10. RNA-seq data analysis

Eight raw RNA-seq files (NCBI SRA FASTQ files) (GEO accession numbers are given in the Table S3) were downloaded onto the Galaxy web server (Galaxy version 24.1.0.dev0)^71^. Of these, n = 4 data files were derived from *Lambdavirus lambda*/*E. coli*-positive placenta samples, and n = 4 data files were derived from *Lambdavirus lambda*/*E. coli*-negative placenta samples. The data were from male placentas (Table S3). Adapters were removed using the Porechop tool, and the reads were further trimmed with the Trim Galore tool to extract high-quality reads with a Phred score ≥10, considered good quality in Nanopore sequencing. Quality-passed reads were mapped to the reference human genome (hg38) using minimap2 (v2.28)^67^. Feature counts were then generated from the BAM files using the featureCounts tool^72^.

Given the small sample size and variations in library size, differential gene expression (DE) analysis between the *Lambdavirus lambda*/*E. coli*-positive and -negative groups was conducted using the weighted limma-voom method with TMM data normalisation and robust settings to minimise biases caused by outliers^28, 29^. The false discovery rate (FDR) was calculated using the Benjamini-Hochberg correction^73^. Differentially expressed genes (DEGs) were identified using an FDR threshold of < 0.05 and a log2 fold-change (log2FC) cutoff of > ±1.5.

Gene set enrichment analysis (GSEA) was performed using the Hallmark gene set database from the Human MSigDB Collections via the WebGestalt 2024 web tool^30, 74^.

### 4.11. H&E and Gram staining of placenta tissue

Snap-frozen villous and decidua basalis tissue samples (1 cm x 1 cm x 2 cm) were fixed in 10% buffered formalin for 48 hours, embedded in paraffin, and sectioned into 5 µm thick slices using the Leica RM2135 microtome. The sections were deparaffinised in Histo-Clear (National Diagnostics, Cat No. HS-200) for 5 minutes twice, then rehydrated through a graded series of industrial methylated spirits (Fisher Scientific, Cat No. M/4400/17) for 5 minutes each, and rinsed in water for 1 minute. The samples were stained with haematoxylin and eosin (H&E) following the manufacturer’s instructions (Sigma-Aldrich, Cat No. 1.05175.0500; National Diagnostics, Cat No. HS-402).

For Gram staining, the hydrated tissue sections were stained with crystal violet solution (Sigma-Aldrich, Cat No. 212525) for 1 minute, followed by rinsing with deionised water. Gram’s iodine solution (Sigma-Aldrich, Cat No. 212527) was then applied for 5 minutes. After washing, the slides were differentiated in alcohol for 10 seconds, rinsed with deionised water, and stained with Safranin O solution (Sigma-Aldrich, Cat No. 212531) for 60 seconds. A final rinse with deionised water was followed by treatment with tartrazine solution (ScyTek, Cat No. TZQ125) for 10 seconds. The slides were dehydrated through a graded series of IMS, cleared with Histo-Clear solution, and mounted with coverslips. Images were captured using a Leica DM3000 microscope at 10x, 20x, 40x air, and 100x oil immersion magnifications.

### 4.12. Statistical analysis

Statistical analyses and graphical presentations were done using GraphPad Prism (v10), the SRplot web server^75^, and the WebGestalt 2024 web tool^74^. The Mann-Whitney U test was used for comparisons between two groups, while one-way ANOVA with Tukey’s post hoc test was applied for comparisons involving more than two groups. A p-value < 0.05 was considered statistically significant. Data are presented as mean ± standard deviation (SD) or as indicated in the respective figure legends.

## Authors contributions

DOCA and KMA conceptualised the study. KMA designed the overall study. MR and TK designed and conducted the metagenomic bioinformatics analyses. NI and KMA developed the placenta tissue sampling protocol. KMA developed the methodology for nucleic acid sample preparations from placenta tissues for shotgun metagenomic sequencing. EA prepared bacterial DNA from CVF samples for NGS sequencing. HH conducted the Nanopore NGS sequencing of placenta and CVF samples. CM and SB performed placental tissue embedding, H&E and Gram staining, imaging, and image analysis. MA and SJ conducted the qPCR array analysis on Bangladeshi placenta samples. KMA performed all major data analyses, including RNA-seq analysis, and prepared the figures and tables. MR, TK, MA, and SJ carried out the initial metagenomic data curation and analysis. KMA wrote the first draft of the manuscript. DOCA, MR, TK, EA, HH, and MA contributed to manuscript writing. NI and AO collected placenta tissue samples at the Cape Town centre. MC provided expertise in placental tissue sampling for metagenomic analysis. MM oversaw patient recruitment in Cape Town. MR oversaw the patient recruitment in Bangladesh. DOCA directed the PRIME project and secured funding for the study.

## Acknowledgements

The authors thank all the members of the PRIME research group at the University of Sheffield, UK, icddr,b Bangladesh and University of Cape Town, South Africa as well as the dedicated midwives, research nurses and doctors at the respective maternity hospitals and clinics for assisting in consenting patients, placenta/CVF collections, tissue harvesting and the Site Files management. We acknowledge Fiona Wright, Histology Technician at the Division of Clinical Medicine and Mark Gell, STH Pathologist for their support in placenta tissue histology and Gram staining, and result interpretation. The study was funded by the United Kingdom’s National Institute for Health Research (NIHR) Global Health Research Scheme to DOCA (GHR 17/63/26). Additional funding by The University of Sheffield Postgraduate Research funds.

## Ethical approvals

This study involving human participants was approved by the London-Fulham Research Ethics Committee, NHS Health Research Authority in the UK (18/LO/2044, Protocol number STH20635); the Faculty of Health Sciences Human Research Ethics Committee (HREC) in Cape Town (HREC REF-196/2019) and the Ethical Review Committee at icddr,b Bangladesh (Protocol number PR-19046).

## Notes

### Competing Interest Statement

The authors have declared no competing interest.

## References

1. Kuperman, A.A. et al. Deep microbial analysis of multiple placentas shows no evidence for a placental microbiome. Bjog 127, 159–169 (2020).

2. Kovalovszki, L., Villányi, Z., Pataki, I., Veszelowvsky, I. & Nagy, Z.B. Isolation of aerobic bacteria from the placenta. Acta Paediatr Acad Sci Hung 23, 357–360 (1982).

3. Stout, M.J. et al. Identification of intracellular bacteria in the basal plate of the human placenta in term and preterm gestations. Am J Obstet Gynecol 208, 226.e221–227 (2013).

4. Aagaard, K. et al. The placenta harbors a unique microbiome. Sci Transl Med 6, 237ra265 (2014).

5. Seferovic, M.D. et al. Visualization of microbes by 16S in situ hybridization in term and preterm placentas without intraamniotic infection. Am J Obstet Gynecol 221, 146.e141–146.e123 (2019).

6. de Goffau, M.C. et al. Human placenta has no microbiome but can contain potential pathogens. Nature 572, 329–334 (2019).

7. Theis, K.R. et al. Does the human placenta delivered at term have a microbiota? Results of cultivation, quantitative real-time PCR, 16S rRNA gene sequencing, and metagenomics. Am J Obstet Gynecol 220, 267.e261–267.e239 (2019).

8. Salter, S.J. et al. Reagent and laboratory contamination can critically impact sequence-based microbiome analyses. BMC Biol 12, 87 (2014).

9. Glassing, A., Dowd, S.E., Galandiuk, S., Davis, B. & Chiodini, R.J. Inherent bacterial DNA contamination of extraction and sequencing reagents may affect interpretation of microbiota in low bacterial biomass samples. Gut Pathog 8, 24 (2016).

10. Chen, C. et al. The microbiota continuum along the female reproductive tract and its relation to uterine-related diseases. Nat Commun 8, 875 (2017).

11. Peric, A., Weiss, J., Vulliemoz, N., Baud, D. & Stojanov, M. Bacterial Colonization of the Female Upper Genital Tract. Int J Mol Sci 20 (2019).

12. Stupak, A. et al. A Virome and Proteomic Analysis of Placental Microbiota in Pregnancies with and without Fetal Growth Restriction. Cells 13 (2024).

13. Manicklal, S., Emery, V.C., Lazzarotto, T., Boppana, S.B. & Gupta, R.K. The “silent” global burden of congenital cytomegalovirus. Clin Microbiol Rev 26, 86–102 (2013).

14. Gindes, L., Teperberg-Oikawa, M., Sherman, D., Pardo, J. & Rahav, G. Congenital cytomegalovirus infection following primary maternal infection in the third trimester. Bjog 115, 830–835 (2008).

15. Gigi, C.E. & Anumba, D.O.C. Parvovirus b19 infection in pregnancy - A review. Eur J Obstet Gynecol Reprod Biol 264, 358–362 (2021).

16. Kirtsman, M. et al. Probable congenital SARS-CoV-2 infection in a neonate born to a woman with active SARS-CoV-2 infection. Cmaj 192, E647–e650 (2020).

17. Njue, A. et al. The Role of Congenital Cytomegalovirus Infection in Adverse Birth Outcomes: A Review of the Potential Mechanisms. Viruses 13 (2020).

18. Radan, A.P. et al. SARS-CoV-2 replicates in the placenta after maternal infection during pregnancy. Front Med (Lausanne) 11, 1439181 (2024).

19. Oliveira, G.M. et al. Detection of cytomegalovirus, herpes virus simplex, and parvovirus b19 in spontaneous abortion placentas. J Matern Fetal Neonatal Med 32, 768–775 (2019).

20. Pesch, M.H., Mowers, J., Huynh, A. & Schleiss, M.R. Intrauterine Fetal Demise, Spontaneous Abortion and Congenital Cytomegalovirus: A Systematic Review of the Incidence and Histopathologic Features. Viruses 16 (2024).

21. Cardenas, I. et al. Viral infection of the placenta leads to fetal inflammation and sensitization to bacterial products predisposing to preterm labor. J Immunol 185, 1248–1257 (2010).

22. Auriti, C. et al. Pregnancy and viral infections: Mechanisms of fetal damage, diagnosis and prevention of neonatal adverse outcomes from cytomegalovirus to SARS-CoV-2 and Zika virus. Biochim Biophys Acta Mol Basis Dis 1867, 166198 (2021).

23. Cruz-Holguín, V.J. et al. Collateral Damage in the Placenta during Viral Infection in Pregnancy: A Possible Mechanism for Vertical Transmission and an Adverse Pregnancy Outcome. Diseases 12 (2024).

24. Schleiss, M.R., Aronow, B.J. & Handwerger, S. Cytomegalovirus infection of human syncytiotrophoblast cells strongly interferes with expression of genes involved in placental differentiation and tissue integrity. Pediatr Res 61, 565–571 (2007).

25. Zuo, T. et al. Gut mucosal virome alterations in ulcerative colitis. Gut 68, 1169–1179 (2019).

26. Reyes, A., Wu, M., McNulty, N.P., Rohwer, F.L. & Gordon, J.I. Gnotobiotic mouse model of phage-bacterial host dynamics in the human gut. Proc Natl Acad Sci U S A 110, 20236–20241 (2013).

27. Akram, K.M., Kulkarni, N.S., Brook, A., Wyles, M.D. & Anumba, D.O.C. Transcriptomic analysis of the human placenta reveals trophoblast dysfunction and augmented Wnt signalling associated with spontaneous preterm birth. Front Cell Dev Biol 10, 987740 (2022).

28. Law, C.W., Chen, Y., Shi, W. & Smyth, G.K. voom: Precision weights unlock linear model analysis tools for RNA-seq read counts. Genome Biol 15, R29 (2014).

29. Liu, R. et al. Why weight? Modelling sample and observational level variability improves power in RNA-seq analyses. Nucleic Acids Res 43, e97 (2015).

30. Liberzon, A. et al. The Molecular Signatures Database (MSigDB) hallmark gene set collection. Cell Syst 1, 417–425 (2015).

31. Caserta, M.T., Hall, C.B., Schnabel, K., Lofthus, G. & McDermott, M.P. Human herpesvirus (HHV)-6 and HHV-7 infections in pregnant women. J Infect Dis 196, 1296–1303 (2007).

32. Ohashi, M. et al. Reactivation of human herpesvirus 6 and 7 in pregnant women. J Med Virol 67, 354–358 (2002).

33. Hendrix, R.W., Smith, M.C., Burns, R.N., Ford, M.E. & Hatfull, G.F. Evolutionary relationships among diverse bacteriophages and prophages: all the world’s a phage. Proc Natl Acad Sci U S A 96, 2192–2197 (1999).

34. Casjens, S.R. & Hendrix, R.W. Bacteriophage lambda: Early pioneer and still relevant. Virology 479-480, 310–330 (2015).

35. Hernandez-Doria, J.D. & Sperandio, V. Bacteriophage Transcription Factor Cro Regulates Virulence Gene Expression in Enterohemorrhagic Escherichia coli. Cell Host Microbe 23, 607–617.e606 (2018).

36. Cools, P. The role of Escherichia coli in reproductive health: state of the art. Res Microbiol 168, 892–901 (2017).

37. Schoch, C.L. et al. NCBI Taxonomy: a comprehensive update on curation, resources and tools. Database (Oxford) 2020 (2020).

38. Davison, A.J. et al. The human cytomegalovirus genome revisited: comparison with the chimpanzee cytomegalovirus genome. J Gen Virol 84, 17–28 (2003).

39. Ross, T.G., Rogers, R.P., Elfrink, N., Bray, N. & Blewett, E.L. Detection of baboon cytomegalovirus (BaCMV) by PCR using primers directed against the glycoprotein B gene. J Virol Methods 125, 119–124 (2005).

40. Tabbuso, T., Defourny, L., Lali, S.E., Pasdermadjian, S. & Gilliaux, O. Moraxella osloensis infection among adults and children: A pediatric case and literature review. Arch Pediatr 28, 348–351 (2021).

41. Hadano, Y. et al. Moraxella osloensis: an unusual cause of central venous catheter infection in a cancer patient. Int J Gen Med 5, 875–877 (2012).

42. Bilyk, V., Ali, O. & Moghrabi, A. [MORAXELLA OSLOENSIS BACTEREMIA WITH PNEUMONIA: FIRST REPORTED CASE IN ISRAEL]. Harefuah 159, 163–165 (2020).

43. Han, X.Y. & Tarrand, J.J. Moraxella osloensis blood and catheter infections during anticancer chemotherapy: clinical and microbiologic studies of 10 cases. Am J Clin Pathol 121, 581–587 (2004).

44. Adapa, S. et al. Peritonitis due to Moraxella Osloensis: An Emerging Pathogen. Case Rep Nephrol 2018, 4968371 (2018).

45. Yamada, A. et al. Peritonitis due to Moraxella osloensis: A case report and literature review. J Infect Chemother 25, 1050–1052 (2019).

46. Li, Y., Wang, G.Q., Ma, X.L. & Li, Y.B. A rare case of acute meningitis caused by Moraxella osloensis. CNS Neurosci Ther 30, e70011 (2024).

47. Gagnard, J.C., Hidri, N., Grillon, A., Jesel, L. & Denes, E. Moraxella osloensis, an emerging pathogen of endocarditis in immunocompromised patients? Swiss Med Wkly 145, w14185 (2015).

48. Lim, J.Y. et al. Complete Genome Sequences of Three Moraxella osloensis Strains Isolated from Human Skin. Genome Announc 6 (2018).

49. LaCroce, S.J. et al. Moraxella nonliquefaciens and M. osloensis Are Important Moraxella Species That Cause Ocular Infections. Microorganisms 7 (2019).

50. Torigoe, K. et al. A Case of Peritoneal Dialysis-Related Peritonitis Due to Moraxella osloensis. Cureus 16, e74294 (2024).

51. Zhang, T., et al. Environmental factors and particle size shape the community structure of airborne total and pathogenic bacteria in a university campus. Front Public Health 12, 1371656 (2024).

52. Jørgensen, N.O., Brandt, K.K., Nybroe, O. & Hansen, M. Delftia lacustris sp. nov., a peptidoglycan-degrading bacterium from fresh water, and emended description of Delftia tsuruhatensis as a peptidoglycan-degrading bacterium. Int J Syst Evol Microbiol 59, 2195–2199 (2009).

53. Shin, S.Y., Choi, J.Y. & Ko, K.S. Four cases of possible human infections with Delftia lacustris. Infection 40, 709–712 (2012).

54. Kawamura, I. et al. Recurrent vascular catheter-related bacteremia caused by Delftia acidovorans with different antimicrobial susceptibility profiles. J Infect Chemother 17, 111–113 (2011).

55. Wiley, L. et al. Bacterial biofilm diversity in contact lens-related disease: emerging role of Achromobacter, Stenotrophomonas, and Delftia. Invest Ophthalmol Vis Sci 53, 3896–3905 (2012).

56. Zhang, L. et al. Characterization and Genome Analysis of the Delftia lacustris Strain LzhVag01 Isolated from Vaginal Discharge. Curr Microbiol 81, 232 (2024).

57. Achermann, Y., Goldstein, E.J., Coenye, T. & Shirtliff, M.E. Propionibacterium acnes: from commensal to opportunistic biofilm-associated implant pathogen. Clin Microbiol Rev 27, 419–440 (2014).

58. Mongaret, C., Velard, F. & Reffuveille, F. Cutibacterium acnes: the Urgent Need To Identify Diagnosis Markers. Infect Immun 89 (2021).

59. Menon, R. et al. Amplification of microbial DNA from bacterial extracellular vesicles from human placenta. Front Microbiol 14, 1213234 (2023).

60. Kaisanlahti, A. et al. Maternal microbiota communicates with the fetus through microbiota-derived extracellular vesicles. Microbiome 11, 249 (2023).

61. Zuo, T. et al. Bacteriophage transfer during faecal microbiota transplantation in Clostridium difficile infection is associated with treatment outcome. Gut 67, 634–643 (2018).

62. Akram, K.M., Frost, L.I. & Anumba, D.O. Impaired autophagy with augmented apoptosis in a Th1/Th2-imbalanced placental micromilieu is associated with spontaneous preterm birth. Front Mol Biosci 9, 897228 (2022).

63. Cavanagh, M. et al. Vaginal host immune-microbiome-metabolite interactions associated with spontaneous preterm birth in a predominantly white cohort. NPJ Biofilms Microbiomes 11, 52 (2025).

64. Stafford, G.P. et al. Spontaneous Preterm Birth Is Associated with Differential Expression of Vaginal Metabolites by Lactobacilli-Dominated Microflora. Front Physiol 8, 615 (2017).

65. Alcolea-Medina, A.a.C., Themoula and Snell, Luke B. and Aydin, Alp and Alder, Christopher and Maloney, Gillian and Bryan, Lisa and Nebbia, Gaia and Douthwaite, Sam and Neil, Stuart and Cliff, Penelope and O’Grady, Justin and Batra, Rahul and Wilks, Mark and O’Hara, Geraldine and Edgeworth, Jonathan. Novel, Rapid Metagenomic Method to Detect Emerging Viral Pathogens Applied to Human Monkeypox Infections. SSRN (2022).

66. De Coster, W., D’Hert, S., Schultz, D.T., Cruts, M. & Van Broeckhoven, C. NanoPack: visualizing and processing long-read sequencing data. Bioinformatics 34, 2666–2669 (2018).

67. Li, H. Minimap2: pairwise alignment for nucleotide sequences. Bioinformatics 34, 3094–3100 (2018).

68. Danecek, P. et al. Twelve years of SAMtools and BCFtools. Gigascience 10 (2021).

69. Wood, D.E., Lu, J. & Langmead, B. Improved metagenomic analysis with Kraken 2. Genome Biol 20, 257 (2019).

70. Lu, J., Breitwieser, F.P., Thielen, P. & Salzberg, S.L. Bracken: estimating species abundance in metagenomics data. PeerJ Comput Sci 3 (2017).

71. Afgan, E. et al. The Galaxy platform for accessible, reproducible and collaborative biomedical analyses: 2018 update. Nucleic Acids Res 46, W537–w544 (2018).

72. Liao, Y., Smyth, G.K. & Shi, W. featureCounts: an efficient general purpose program for assigning sequence reads to genomic features. Bioinformatics 30, 923–930 (2014).

73. Robinson, M.D., McCarthy, D.J. & Smyth, G.K. edgeR: a Bioconductor package for differential expression analysis of digital gene expression data. Bioinformatics 26, 139–140 (2010).

74. Elizarraras, J.M. et al. WebGestalt 2024: faster gene set analysis and new support for metabolomics and multi-omics. Nucleic Acids Res 52, W415–w421 (2024).

75. Tang, D. et al. SRplot: A free online platform for data visualization and graphing. PLoS One 18, e0294236 (2023).

