## Supplementary material for "Distinct Virome and Bacteriome Profiles of Term and Preterm Placentas from African, Asian and European Women": Akram et al Supplementary file.docx

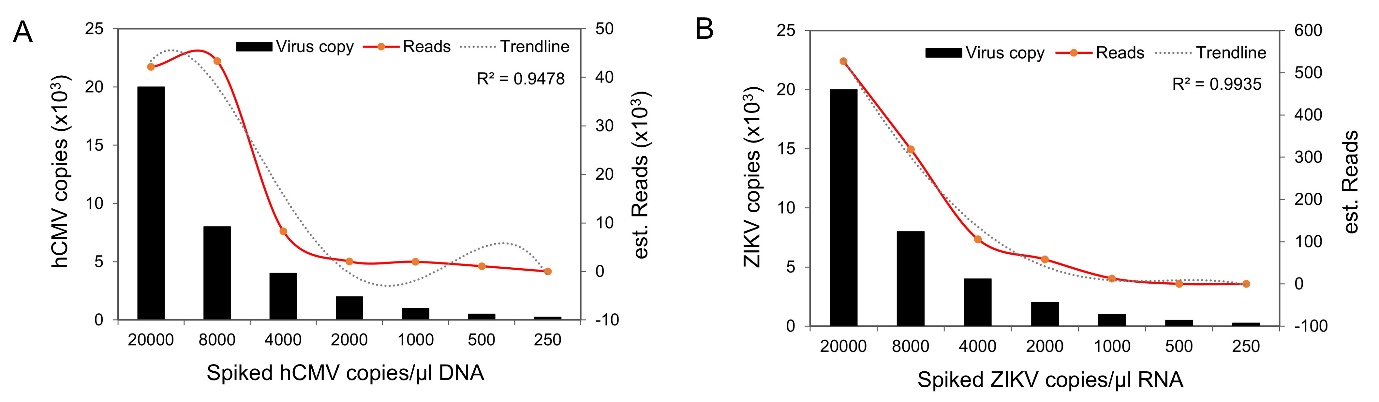

**Figure S1:** Viral nucleic acids spiking doses versus read counts graphs showing detection of hCMV (A) and ZIKV (B) in placenta samples by shotgun sequencing. X-axis represents nucleic acids spiking concentrations, left Y-axis represents the estimated viral copies and right Y-axis represents transcript counts by sequencing.

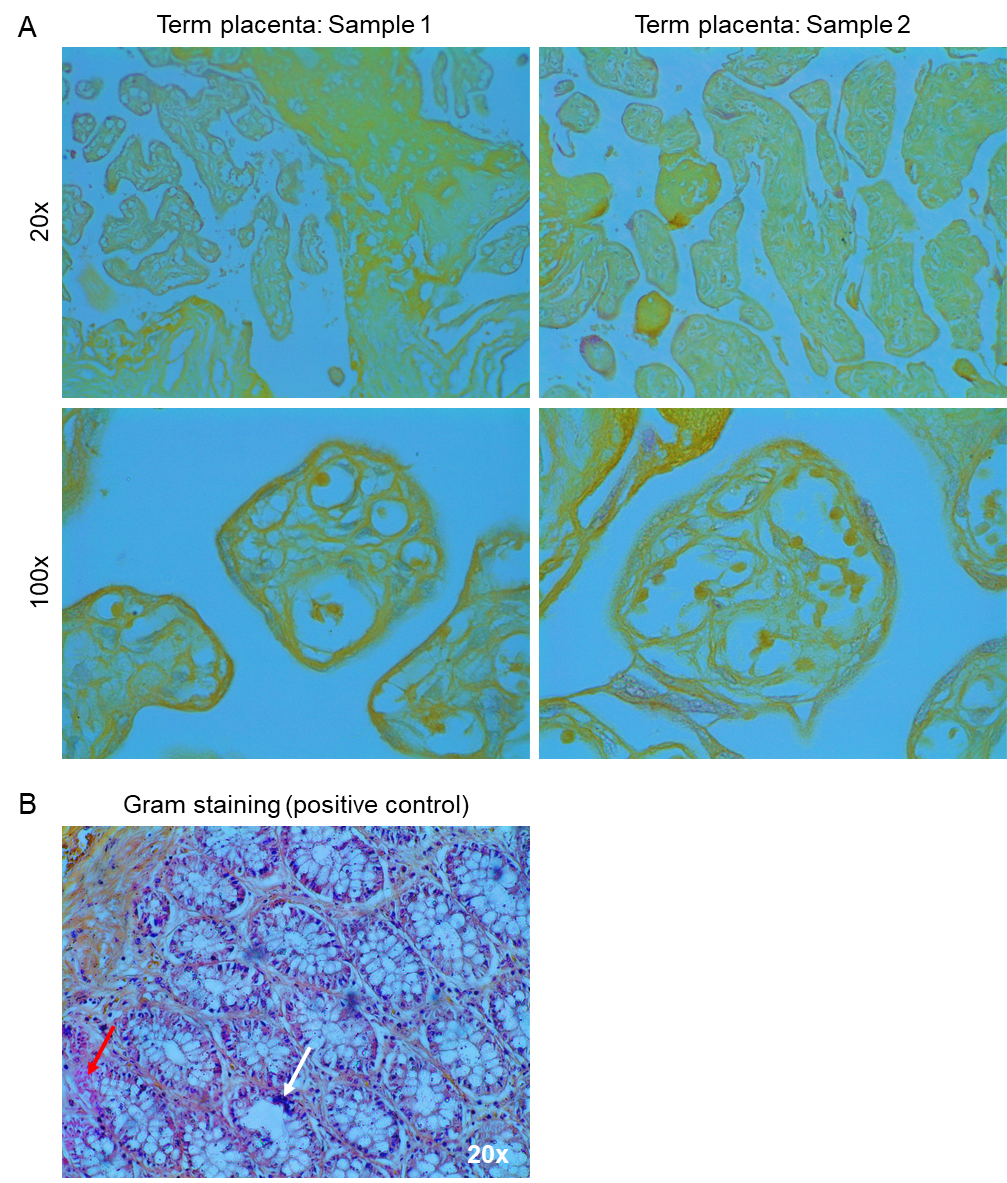

**Figure S2:** Gram staining on FFPE term sequence-negative placenta villous/decidua tissue section. A. Representative microscopic images (20x and 100x oil immersion) from two independent term placenta samples showing negative staining for Gram staining. B. Positive control for Gram staining showing both gram-positive (purple colour, white arrow) and gram-negative (pink colour, red arrow) bacteria within the tissue.

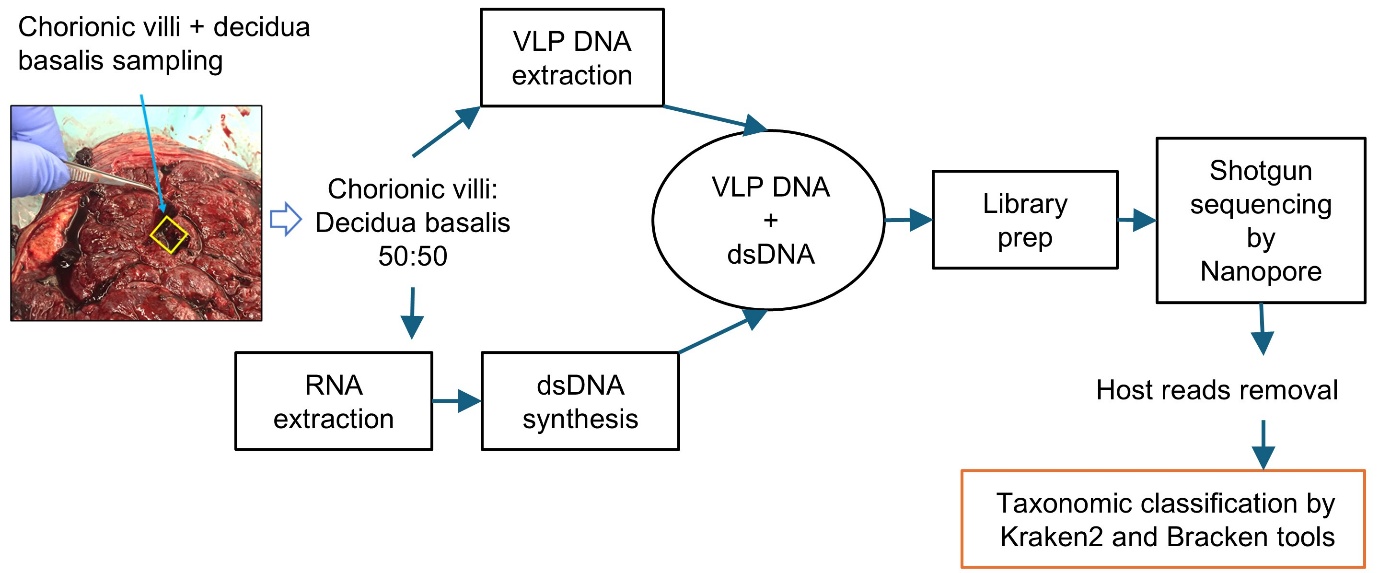

**Figure S3:** Flowchart showing metagenomic sequencing and data analysis pipeline.
